# Evolutionary origins and transcriptomic innovations of vertebrate Cajal-Retzius cells

**DOI:** 10.1101/2025.08.21.671508

**Authors:** Elias Gumnit, Giacomo Gattoni, Jamie Woych, Astrid Deryckere, Prabesh Bhattarai, J Andrew Gillis, Caghan Kizil, Maria Antonietta Tosches

## Abstract

Evolutionary changes of cortical development were instrumental for the emergence of the mammalian neocortex. Cajal-Retzius cells, a transient neuron type discovered over a century ago, are critical players in the development of mammalian-specific cortical features such as inside-out corticogenesis. However, it is unclear whether Cajal-Retzius cells exist only in mammals, or if they are ancestral in vertebrates but acquired new functions during mammalian brain development. To trace the evolution of this cell type, we probe the presence of Cajal-Retzius cells in chicken, salamander, zebrafish, and little skate. First, through comparative transcriptomics and spatial analysis we show that Cajal-Retzius cells are conserved in vertebrates. However, only amniote Cajal-Retzius cells gained robust expression of *Reelin*, a secreted glycoprotein crucial for mammalian cortical development. Second, we find that Cajal-Retzius cells are part of a larger and diverse family of neuron types, and are closely related to *Tp73*+ external tufted cells in the olfactory bulb, which express most of the “canonical” Cajal-Retzius transcription factors. Our results indicate that Cajal-Retzius cells emerged early in vertebrate history, using a regulatory program largely shared with neurons in the olfactory system, and that transcriptomic novelties underlie their specialized roles in mammalian cortical development. This is an extreme example of how cell types may diverge in morphology and function over the course of evolution even when their core transcriptomic profile is broadly preserved.

## Introduction

The forebrain is the brain region that underwent the most dramatic changes in vertebrate evolution ^1^. In mammalian ancestors, the drastic expansion of the pallium (dorsal telencephalon) occurred with several innovations, including the anatomical and functional diversification of cortical regions, a six-layered neocortex, and a folded hippocampus. Changes in cortical development may underlie the evolution of these novelties ^1,2^. To investigate the underpinnings of these changes, we focused on Cajal-Retzius (CR) cells, a multifunctional cell type with critical roles in the development of mammalian-specific cortical traits ^3–6^. In mammals, CR cells are born in the ventral pallium, cortical hem, septum, and prethalamic eminence (PThE), and then migrate tangentially in the marginal zone to cover the entire telencephalic vesicle ^7–12^. These cells are the major source of Reelin, a secreted molecule that regulates the radial migration of immature neurons during corticogenesis ^13,14^. CR cells also control cortical arealization, serve as pioneer neurons in the hippocampus and olfactory system, influence hippocampal morphogenesis, and regulate circuit assembly in the hippocampus and neocortex ^8,14–25^. Clarifying the evolution of CR cells is thus essential to understand how innovations in the mammalian cerebral cortex emerged.

Notably, while CR cells are the first neurons that develop in the mammalian pallium, their evolution remains highly debated. Markers of mammalian CR cells, including Reelin and the transcription factor p73, are expressed in scattered pallial cells in developing lizard, crocodile, chicken, frog, and fish brains ^26–30^. A recent study proposed that CR cells existed already in tetrapod ancestors ^30^, but the interpretation of chick data is contentious ^29,31,32^, and fishes remain poorly investigated. The expression of single CR markers remains insufficient to conclude whether CR cells exist in non-mammals: none of these markers are CR-specific, but it is only their *combined* expression that defines CR identity^3^.

With the advent of single-cell RNA sequencing, mammalian CR cells have been well characterized at the transcriptomic level, and express a core set of transcription factors (TFs) including: *Tp73, Tbr1, Lhx1, Lhx5, Ebf2, Ebf3, Nr2f2, Nhlh2,* and *Barhl2* (with *Tp73* and *Ebf3* missing in ventral-pallium derived cells) ^5,33^. Among these, *Tp73* (*Trp73* in mice) encodes for p73 and is expressed in CR cells derived from medial sources (PThE, septum, and hem). (For simplicity, hereafter we refer to the mouse *Trp73* gene as *Tp73*). *Tp73* is a key player in the gene regulatory network that establishes medial CR identity: mutant mice display abnormal cortical lamination and hippocampal malformations in conjunction with loss of medial CR cells ^26,29^.

Here we explored the evolution of CR cells by examining species that represent the main branches of the vertebrate tree. Expanding on previous work^27–30,34^, we found *Tp73*+ cells in the pallium and other conserved spatial locations in all vertebrate species examined. Motivated by these observations, we then performed unbiased transcriptomic comparisons to identify CR-like neurons in non-mammalian species. In-depth analyses revealed that *Tp73* defines a diverse and evolutionarily-conserved family of neurons, including not only CR cells, but also neurons in the olfactory bulb (OB) and in the basal forebrain. With our cross-species analyses, we identify the conserved set of transcription factors that differentiate CR cells from their closely-related neuron types, and identify critical evolutionary innovations of the CR cell transcriptome. Taken together, our findings indicate that CR cells were already present in vertebrate ancestors and share evolutionary origins with a neuron type in the OB, raising new scenarios on the early evolution of this enigmatic cell type.

## Results

### Conserved populations of *Tp73*+ neurons in vertebrates

In the developing mouse brain, *Tp73* is expressed in neurons distributed within the pallium, subpallium, olfactory bulb (OB), preoptic area (POA), and prethalamic eminence (PThE, a diencephalic region that persists after development only in amphibians and fish)^35^. *Tp73* is expressed also in some non-neuronal cells including the choroid plexus, which develops from the cortical hem (Fig. 1A) ^36^. Pallial *Tp73+* neurons are CR cells and express *Reln*, whereas diencephalic *Tp73*+ neurons that do not coexpress *Reln* are not classified as CR cells ^35,37^ (Fig. 1A).

**Figure 1.**
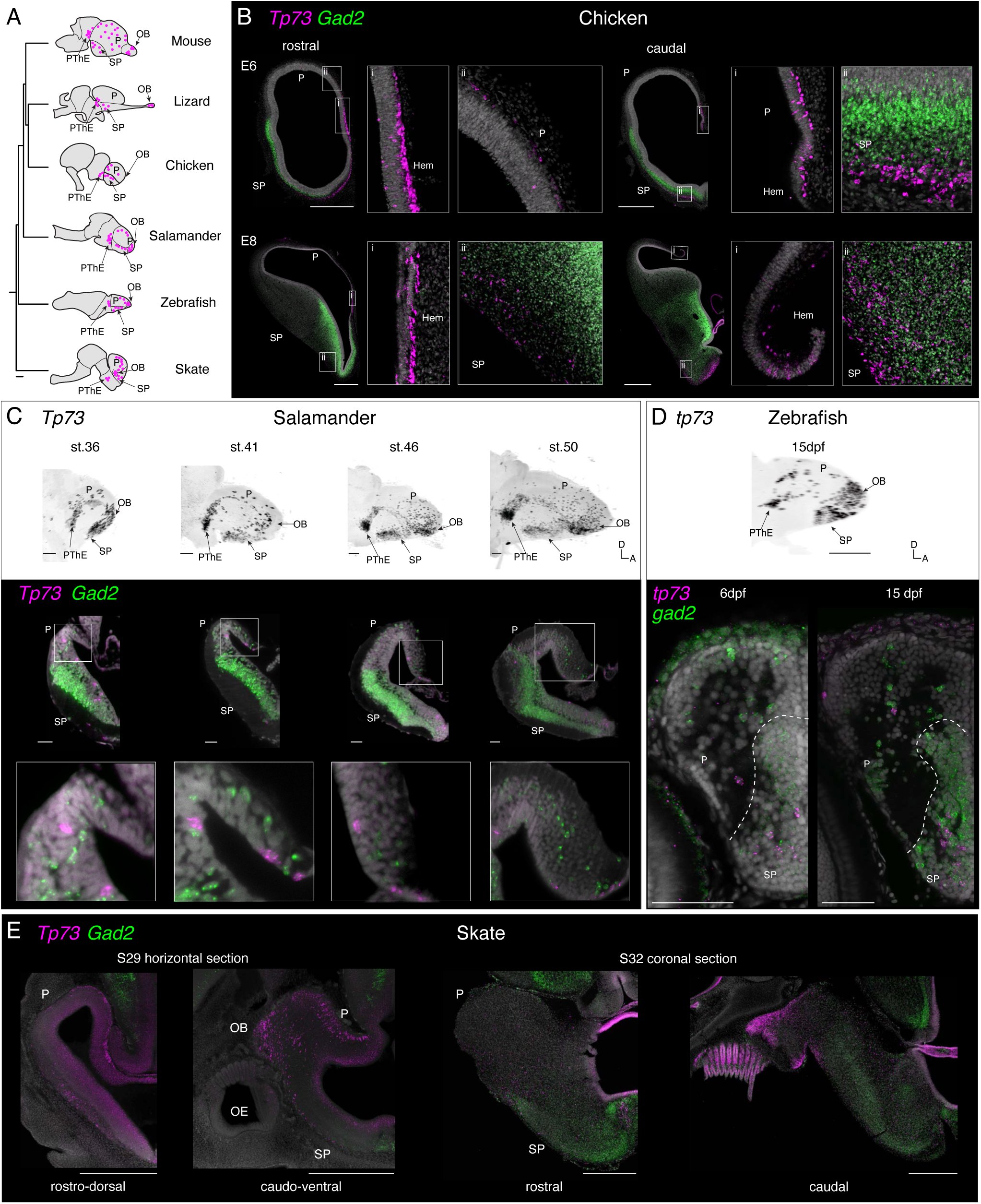
Conserved *Tp73* expression in vertebrate forebrains. (A) Distribution of *Tp73* neurons in vertebrate brains, according to published data and this study. Scale bar: 100 million years. (B) Double HCRs for *Tp73* (magenta) and *Gad2* (green) in coronal sections of chick brains (embryonic stages E6 and E8), showing *Tp73* expression in the developing hem, pallium, and subpallium. *Gad2* labels GABAergic neurons and helps identify the pallial-subpallial boundary. Scale bars = 500 um. (C) Top: 3D max intensity projections (lateral views, anterior on the right) of salamander brains (larval stages 36, 41, 46, and 50) after whole mount HCR for *Tp73.* Scale bar 100um. Bottom: optical coronal sections through one telencephalic hemisphere of *Tp73* and *Gad2*-labeled brains at different developmental stages; insets show *Tp73*+ and *Gad2*+ neurons in pallium. Scale bar 50 um. (D) 3D max intensity projection of a whole mount 15 dpf zebrafish brain (lateral view, anterior on the right) after HCR for *tp73.* Scale bar 100 um. Bottom: sections of *tp73* and *gad2*-labeled 6 dpf and 15dpf zebrafish brains showing *tp73+* and *gad2*+ neurons in pallium and subpallium. Scale bar 50 um. (E) Expression of *Tp73* and *Gad2* in the developing skate brain. Left: horizontal sections of stage 29 brains; right: coronal sections of stage 32 brains. Scale bar 500um. See additional sections in fig. S1. Abbreviations: E, embryonic day; P, pallium; SP, subpallium; OB, olfactory bulb; OE, olfactory epithelium; PThE, prethalamic eminence; dpf, days post fertilization; A, anterior; D, dorsal.

Using HCR *in situ* hybridization, we characterized the distribution of *Tp73*+ neurons during brain development in multiple species representing the major vertebrate clades. Scattered *Tp73*+ neurons have been reported in the developing chick brain ^29^. These *Tp73*+ neurons are most dense near the cortical hem and grow sparser laterally into the medial and dorsal pallium (Fig. 1B). We also observed *Tp73+* neurons in the chick subpallium, PThE, and OB (Fig. 1B, fig. S1A). Overall, the distribution pattern was similar to the one previously observed in lizard embryos, where *Tp73+* neurons were found in the PThE, subpallium, and OB, plus sparse *Tp73+* neurons in the pallium ^27^ (Fig. 1A). In the salamander *Pleurodeles waltl,* we found scattered *Tp73*+ neurons also distributed throughout the pallium - with higher density around the hem - from early to late stages of pallial neurogenesis ^38^. In addition, *Tp73*+ cells were visible in the OB, PThE, and subpallium (Fig. 1C, fig. S1B, video S1).

We then extended our analysis to two distantly-related fish species: the teleost zebrafish and a cartilaginous fish, the little skate *Leucoraja erinacea*. In the developing zebrafish forebrain, *tp73*+ cells were present in the same brain regions described in tetrapods, with sparse *tp73*+ neurons in the pallium (Fig. 1D, fig S1C, video S2). In the developing skate, *Tp73*+ neurons were found in high density in the OB and pallium, and more sparsely in the subpallium and putative PThE, consistent with other vertebrates (Fig. 1E, fig. S1D).

In the adult mouse brain, *Tp73*+ neurons persist in the OB, basal forebrain, and pallium, although only a small percentage of CR cells in the pallium survives to adulthood ^39^. Consistently, we found *Tp73*+ neurons in the same brain areas, plus the PThE, in adult salamander, zebrafish, little skate, and in the bichir fish *Polypterus senegalus* (a non-teleost ray-finned fish), but with a drastic reduction of the density of *Tp73*+ pallial cells, particularly in skates (fig. S2, video S3-4). These results indicate that the spatial pattern of *Tp73* expression is conserved across jawed vertebrates, with at least some *Tp73* neurons in the developing pallium, where in mammals *Tp73* is highly specific to CR cells born from medial sources (cortical hem, septum, and PThE).

Motivated by these observations, we asked whether neurons resembling mammalian CR cells at the transcriptome level exist in the developing forebrain of non-mammalian species. For this we relied on our recent high quality scRNA-seq dataset of the developing salamander brain^40^, which spans eight developmental stages and multiple brain regions. Only one cluster in this dataset expressed high levels of *Tp73* (Fig. 2A-B). This cluster also expressed neuronal markers, such as the synaptic gene *Snap25* and the vesicular glutamate transporter gene *Slc17a6*, which clearly differentiated it from non-neuronal cells expressing low levels of *Tp73* (fig. S3B). Importantly, TFs usually associated with mammalian CR cells, including *Tp73*, *Lhx1*, *Lhx5*, *Nr2f2*, *Ebf2*, *Ebf3*, and *Barhl2* ^5^, were co-expressed only in the *Tp73*+ neuronal cluster (Fig. 2B, fig. S3A). To further determine whether these salamander *Tp73*+ neurons are the most similar to mouse CR cells at the transcriptomic level, we compared our scRNAseq data with a developmental mouse brain dataset ^41^, restricting our analysis to mature neurons (see Methods). We used three independent methods to compare the salamander and mouse datasets. First, we computed all the genes that differentiate mouse CR cells from other mouse differentiated neurons, and then assessed the relative expression enrichment for these genes in the salamander dataset. *Tp73* neurons in salamander showed the highest enrichment for this CR gene module (Fig. 2C). Second, we computed pairwise transcriptomic correlations (gene specificity index ^42^, see Methods) of all salamander and mouse neuron classes, and found that the salamander *Tp73*+ neurons are most similar to mouse CR cells (Fig. 2D). Third, using manifold integration ^43^ for the joint analysis of the two data sets, we identified a cluster in the integrated transcriptomic space that includes salamander *Tp73*+ neurons and mouse CR cells, supporting the transcriptomic similarity of these cell types (Fig. 2E, fig. S4A-D). These data suggest a close transcriptomic similarity of salamander *Tp73*+ neurons and mouse CR cells.

**Figure 2.**
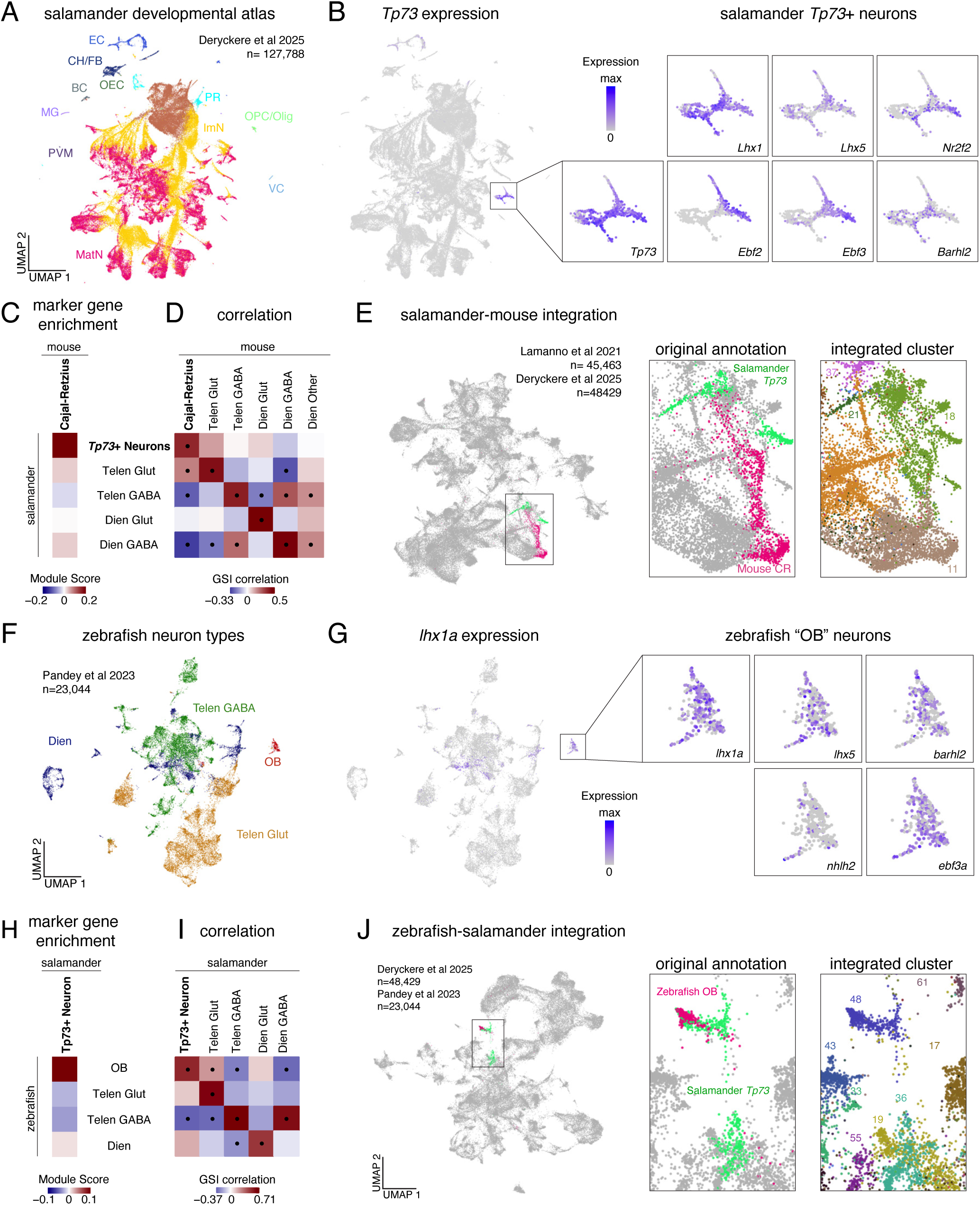
Mouse, salamander, and zebrafish *Tp73*+ neurons have a shared transcriptomic signature. (A) UMAP of the Deryckere et al. ^40^ scRNA-seq dataset from the developing salamander brain (n=127,788 cells), showing cells colored by broad class. (B) Left: UMAP showing *Tp73* expression in the developing salamander brain dataset. Right: Zoom on *Tp73+* neurons, showing the expression of known CR marker TFs ^5^ (fig. S3A for full UMAPs of zoomed genes). (C) Expression enrichment of mouse CR cell markers (Seurat Module Score) in salamander cell classes. Module score computed for the top 49 CR markers in La Manno et al ^41^ (see Methods for gene selection criteria). (D) Pairwise GSI correlations of classes of salamander and mouse forebrain neurons (mouse data from La Manno et al.^41^). Correlations based on the intersection of genes that were differentially expressed in both mouse and salamander (336 total genes). Dots indicate statistically significant correlations. (E) UMAP showing forebrain neurons from developing salamander and mouse after Seurat CCA integration. Salamander *Tp73*+ neurons and mouse CR cells highlighted in green and magenta, respectively. Right: Zoom on integrated cluster 18 containing salamander *Tp73* neurons and mouse CR cells. (F) UMAP of the Pandey et al 2023 ^44^ dataset from the developing zebrafish telencephalon (n=23,044 cells, stages 6 and 15 dpf only), showing cells colored by class. (G) Left: UMAP showing *lhx1a* expression in the larval zebrafish dataset. Right: Zoom on zebrafish OB neurons showing expression of *lhx1a* and other CR TFs. (H) Expression enrichment of salamander *Tp73*+ neurons markers (Seurat Module Score) in zebrafish cell classes. Module score computed for the top 42 markers of salamander *Tp73+* neurons (see methods for gene selection criteria) (I) Pairwise GSI correlations of salamander and zebrafish forebrain neuron classes. Correlations based on the intersection of genes that were differentially expressed in both zebrafish and salamander (143 total genes). Dots indicate statistically significant correlations. (J) UMAP showing forebrain neurons from developing salamander and zebrafish after Seurat CCA integration. Right: Zoom on integrated cluster 48 containing salamander *Tp73* neurons and zebrafish OB cells. Abbreviations: BC, blood cells; CH/FB, chondrocytes/fibroblasts; EC, epithelial cells; Dien, diencephalon; GABA, GABAergic; Glut, glutamatergic; GSI, gene specificity index; ImN, immature neurons; MatN, mature neurons; MG, microglia; OEC, olfactory ensheathing cells; OPC/Olig, oligodendrocyte precursor cells/oligodendrocytes; PVM, perivascular macrophages; PR, photoreceptor cells; RG, radial glia; Telen, telencephalon; VC, vascular cells; see also Fig. 1.

Next, we analyzed transcriptomic data from zebrafish to assess the existence of neurons similar to salamander *Tp73*+ neurons. In a dataset by Pandey et al. ^44^ from the developing zebrafish forebrain, a glutamatergic cell type annotated as “olfactory bulb” (OB) expressed many of the TFs characterizing mammalian CR cells and salamander *Tp73*+ neurons such as *lhx1a*, *lhx5*, *ebf3*, and *barhl2* (Fig. 2F-G, fig. S5A-F). *tp73* was poorly detected in this dataset, probably due to technical reasons; supporting this conclusion, this gene was detected in a more deeply-sequenced adult dataset ^45^, where it was coexpressed with other CR TFs (see fig. S5L-S). The OB cluster also showed the highest enrichment score for genes that distinguish *Tp73*+ neurons from other neuron types (Fig. 2H). GSI correlation analysis of zebrafish and salamander neurons showed a significant correlation between salamander *Tp73*+ neurons and the zebrafish OB cluster. (Fig. 2I). In addition, integration of salamander and zebrafish data showed co-clustering of zebrafish OB cells and a subset of salamander *Tp73*+ neurons; a second group of salamander *Tp73*+ neurons co-clustered with zebrafish diencephalic neurons, suggesting transcriptomic heterogeneity (Fig. 2J, fig. S4E-H). We conclude that the zebrafish OB cluster has high similarity to salamander *Tp73*+ neurons (markers of OB mitral cells, the major OB glutamatergic type, are expressed in a separate set of cells, see fig. S5H-K). Taken together, these results indicate that *Tp73*+ neurons are overall highly similar at the transcriptomic level in mouse, salamander, and zebrafish, supporting their evolutionary conservation in vertebrates.

### Heterogeneity of salamander *Tp73*+ neurons

The presence of *Tp73*+ cells in different areas of the forebrain (Fig. 1), together with the varied expression of marker genes in *Tp73*+ neurons (Fig. 2) suggests they are heterogeneous. Therefore, we sought to further characterize the molecular heterogeneity of salamander *Tp73*+ neurons in order to clarify their comparison with mammalian CR cells. To increase the power of our analyses, we merged *Tp73*+ clusters from larval and adult ^46^ scRNA-seq datasets, after noticing that *Tp73*+ neurons have conserved transcriptomes and spatial distribution across the salamander lifespan (fig. S2A-B). This salamander *Tp73*+ dataset, restricted to neurons, contained 982 cells (Fig. 3A, fig. S6).

**Figure 3.**
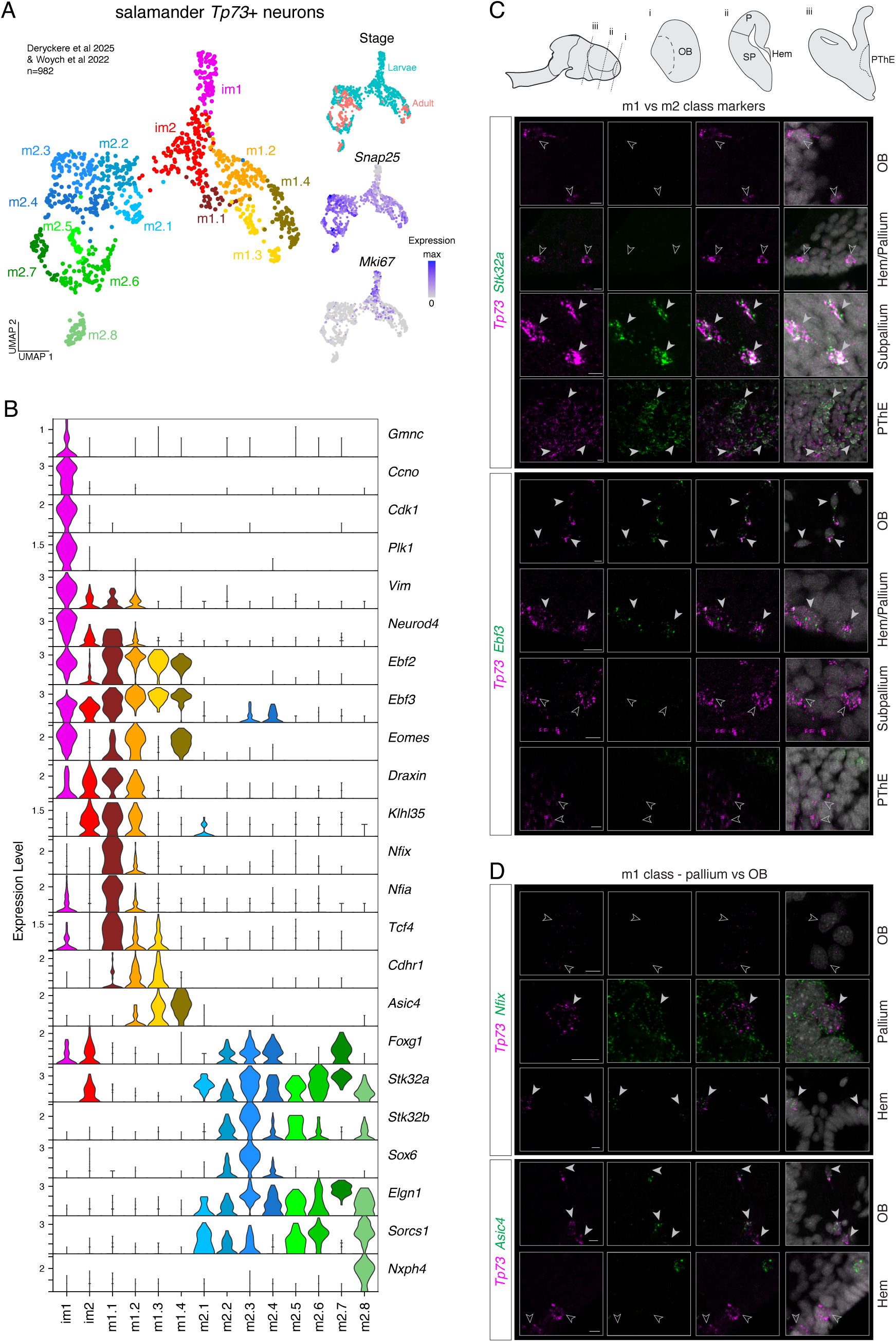
Transcriptomic and spatial heterogeneity of salamander *Tp73+* neurons. (A) UMAPs of the salamander *Tp73*+ neurons dataset (n= 982 cells). Left: *Tp73*+ neurons colored by cluster. Right: *Tp73*+ neurons colored by dataset (adult from Woych et al ^46^), and expression of *Snap25* and *Mki67*. (B) Violin plots showing the expression of marker genes for *Tp73* neuronal clusters; m1 and m2 populations express different sets of markers. (C)-(D) Closeups showing coexpression of *Tp73* and selected marker genes detected by HCR in late active larvae (stages 46-55; see fig S8 for full images). (C) Top: coexpression of *Stk32a* and *Tp73* in PthE and subpallium but not in OB and hem/pallium. Bottom: *Ebf3* and *Tp73* coexpression in OB and hem/pallium but not subpallium and PThEregions. Scale bar 10um. (D) Top: coexpression of *Nfix* and *Tp73* in pallium but not OB. Bottom: coexpression of *Asic4* and *Tp73* in OB but not hem. Scale bar: 10 um. Abbreviations: im, immature; m, mature; see also Figure 1.

After clustering and differential gene expression analysis, we identified three major classes of *Tp73*+ neurons, each including multiple clusters. The first class comprised late progenitors and immature neurons, as indicated by the expression of the proliferation marker *Mki67*, low levels of the synaptic gene *Snap25*, and markers of immature neurons like *Vim* and *Neurod4* (clusters im1-2, Fig. 3A-B). These clusters were uniquely labeled by *Gmnc* and *Cccno*, members of a multiciliation gene module expressed transiently during early CR differentiation in mice ^29^. Double HCR *in situ* hybridization showed coexpression of *Tp73* with *Gmnc* and *Ccno* in sparse cells in the hem, septum, and PThE (fig. S7). These observations suggest that cells in clusters im1-2 are in the process of acquiring a terminal CR-like neuronal fate.

The other two classes were identified as mature *Tp73*+ neurons. The first class (clusters m1.1 to 1.4) shared some of the defining features of CR cells, including *Ebf2* and *Ebf3* expression and absence of *Foxg1* expression. The second class (clusters m2.1 to m2.8) expressed the unique combination of *Stk32a*, *Elfn1*, and *Sorcs1* (Fig. 3B). We performed HCRs to locate these two distinct populations in the developing brain. *Stk32a* and *Tp73* (m2 class) were coexpressed in the subpallium and PThE, but not in the OB or pallium. In contrast, *Ebf3* and *Tp73* (m1 class) were coexpressed in cells in the OB and pallium, but not in the subpallium or PThE (Fig. 3C, fig. S8A-B). We conclude that mature salamander *Tp73*+ neurons belong to spatially and molecularly distinct populations: a population of *Tp73*+ *Ebf3*+ neurons enriched in the anterior and dorsal forebrain, and a second population of *Tp73*+ *Ebf3*-neurons in the posterior and ventral forebrain. These two neuronal classes co-cluster with different sets of neurons in the zebrafish dataset, suggesting an ancient origin (fig. S8C).

Next, we explored further the heterogeneity of *Tp73*+ *Ebf3*+ neurons. Cluster m1.1 expressed high levels of *Nfix* and *Nfia*, markers of mammalian hem-derived CR cells ^33^. When staining for *Nfix*, we found coexpression with *Tp73* only in the pallium, including the hem region, and not in the OB (Fig. 3D, fig S8D-E). Conversely, clusters m1.2-4 expressed *Asic4* and *Cdhr1*, markers of mammalian OB glutamatergic neurons ^47,48^. *Asic4* showed the opposite pattern as *Nfix*, as it was coexpressed with *Tp73* in OB but not in pallium (Fig. 3D, fig S8D). This suggests that the *Tp73*+ *Ebf3*+ class of salamander neurons includes a pallial population with additional markers of mammalian CR identity, and an OB population expressing markers of OB glutamatergic neurons.

### Cajal-Retzius and olfactory bulb *Tp73*+ external tufted cells are sister cell types

Next, we investigated whether the molecular and spatial heterogeneity of salamander *Tp73*+ neurons has counterparts in mammals. We isolated mouse *Tp73+* neurons from the Allen Brain Cell (ABC) atlas ^39^ and compared them to the salamander *Tp73*+ neuron dataset. The ABC atlas, comprising the entire adult mouse brain, was chosen for its high sequencing depth, completeness, and matching MERFISH data, necessary to resolve subtle differences between *Tp73*+ neurons types. This atlas includes three populations of *Tp73+* neurons (Fig. 4A and fig. S9A-B). One is composed of subclass 118 ADP-MPO Trp73 and cluster 2111 LHA Barhl2 Glut_2, spatially located in the basal forebrain, specifically in areas annotated as the lateral hypothalamus and medial POA. A second group is the subclass 036 HPF CR Glut, which includes CR cells sampled from the adult cortex and hippocampus (Fig. 4A, fig. S9A). Finally, a third group is the supertype 0132 OB Eomes Ms4a15 Glut_3, which was further characterized by Zeppilli et al ^47^ in a transcriptomic analysis of the OB. These cells express *Lhx1* and *Ebf3* and correspond to one type of external tufted cells (ETCs) (see cluster ET2 in Zeppilli et al ^47^). ETCs are a heterogeneous class of glutamatergic neurons in the mammalian OB; these cells, or at least a subset of them, are believed to be born in the pallium before migrating to the bulb ^49–51^. The *Tp73+* ETCs are juxtaglomerular tufted cells that synapse with olfactory sensory neurons and other periglomerular interneurons ^52,53^ (fig. S10). Using cross-species data integration, we found that classes of salamander *Tp73*+ neurons integrate with distinct mouse *Tp73*+ neurons. We computed the overlap of cells’ cross-species neighborhoods in the integrated transcriptomic space (see Methods). Salamander *Tp73*+ *Ebf3*-neurons were on average more similar to mouse basal forebrain *Tp73*+ neurons, while salamander *Tp73*+ *Ebf3*+ neurons were more similar to mouse CR and ETC (Fig. 4C). Specifically, salamander clusters m1.1 (*Nfix+*) and m1.2 were most likely to be mutual nearest neighbors with mouse CR cells (Fig. 4C) and also co-clustered with them (fig. S11A-C), while m1.4 (*Asic4+*) neurons were nearest neighbors with mouse ETC cells in the integrated space (Fig. 4C) and co-clustered together (fig.S11A-C). GSI correlation analysis showed a similar trend, although this analysis was less sensitive than integration at distinguishing CR and ETC cells due to the high transcriptomic similarity between these two cell types (Fig. 4D). These comparisons indicate that salamander *Tp73*+ *Ebf3*+ *Nfix*+ neurons are homologous to mammalian CR cells, while salamander *Tp73*+ *Ebf3*+ *Asic4*+ neurons are homologous to ETCs.

**Figure 4.**
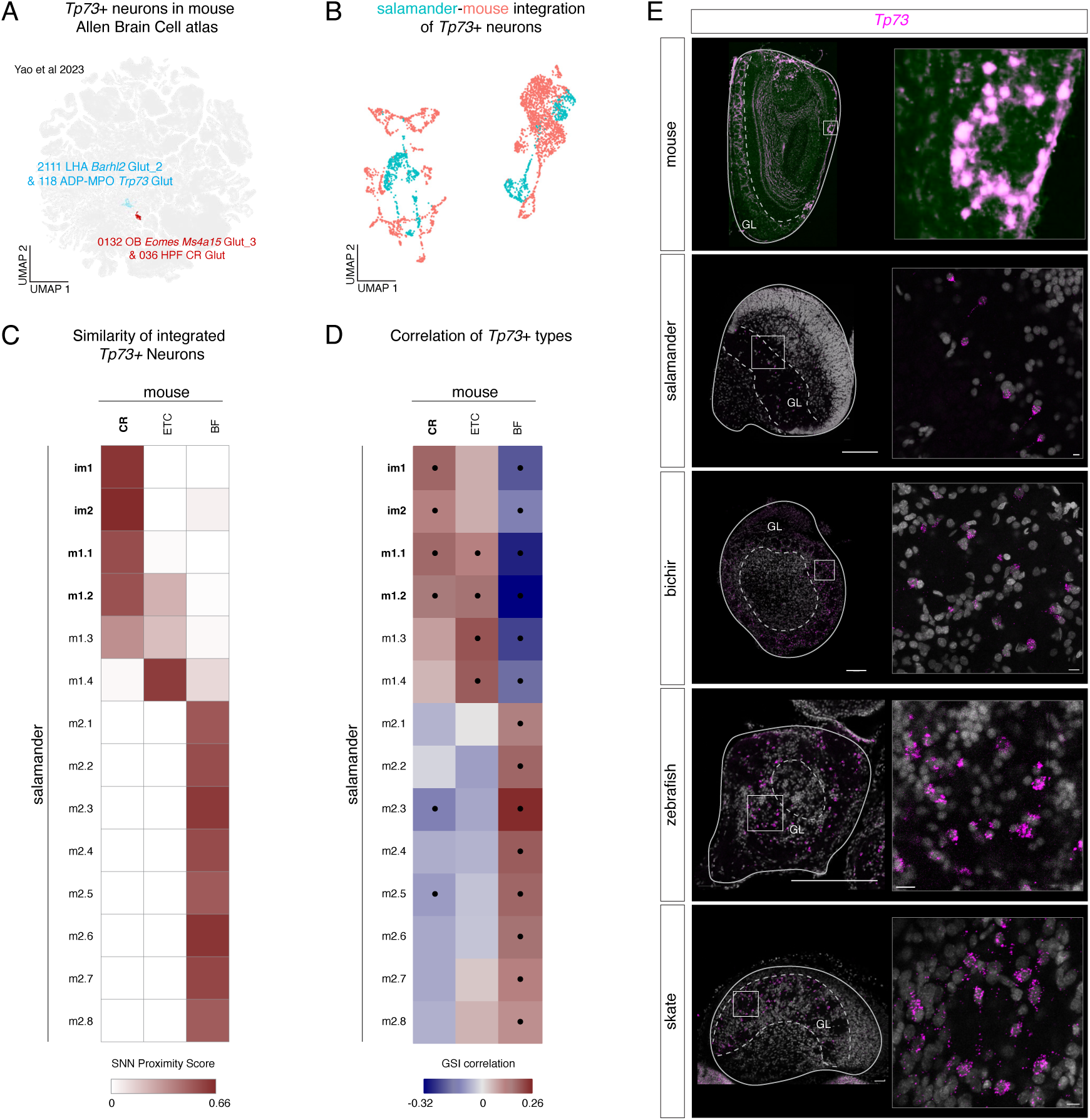
Transcriptomic similarity of CR and *Tp73*+ external tufted cells in mouse and salamander. (A) UMAP of the mouse Allen Brain Cell atlas ^39^, highlighting the *Tp73*+ cells used for further analysis. (B) UMAPs showing salamander and mouse *Tp73+* neurons after Seurat CCA integration, with cells colored by species. (C) Cross-species nearest neighbor proximity of mouse and salamander *Tp73*+ neurons in the integrated transcriptomic space. Higher score indicates higher reciprocal proximity of neurons of a given type in the integrated space (see Methods). (D) Pairwise GSI correlations of mouse and salamander *Tp73*+ types. Correlations based on the intersection of genes differentially expressed in both mouse and salamander (647 total). Dots indicate statistically significant correlations. (E).*Tp73*+ neurons in the OB glomerular layer of mouse (adult), salamander (adult), bichir fish (adult), zebrafish (3 mpf), and skate (adult). Mouse *Trp73* colorimetric *in situ* hybridization adapted from mouse Allen Mouse Brain Atlas, mouse.brain-map.org and atlas.brain-map.org ^48^. All other images are *Tp73* HCRs. Full image scale bars 200um. Zoom scale bars 10um. Abbreviations: BF, basal forebrain; ETC, external tufted cells; GL, glomerular layer; see also Figures 1, 2, and 3.

Next, we wondered whether transcriptomically distinct CR and ETC cells exist in other vertebrate species. The available chicken ^54^ and zebrafish ^44^ datasets had several limitations, including low numbers of *Tp73*+ neurons, absence of OB sampling (in chicken), and relatively lower sequencing depth (in zebrafish). Nevertheless, comparisons of chicken and zebrafish *Tp73*+ neurons with mouse ABC Atlas *Tp73*+ neurons supported the conclusion that the chicken *Tp73*+ neurons in Zaremba et al. ^54^ are CR cells (fig. S10, fig. S11D-F), and that most zebrafish *tp73*+ neurons in Pandey et al. ^44^ are ETCs (fig. S11G-I). The fact that a few zebrafish cells co-clustered with mouse CR cells (fig. S11G-I), together with our tissue staining data (Fig. 1), is consistent with the possibility that CR cells also exist in teleosts.

To further support the existence of *Tp73*+ ETCs across vertebrates, we analyzed *Tp73* expression in the mature OB, where glomeruli are well defined. The mouse *Tp73+* ETCs have a characteristic spatial distribution: they surround the olfactory glomeruli ^47^ (Fig. 4G). We observed *Tp73*+ neurons in the glomerular layer of salamander, bichir, zebrafish, and skate (Fig. 4G). In the three fish species, *Tp73* neurons were arranged in characteristic rings around the glomeruli (Fig. 4G). These results indicate that *Tp73*+ ETCs are a conserved neuron type in jawed vertebrates.

Finally, we asked whether the expression of similar sets of transcription factors underlies the transcriptomic similarity of CR cells and ETCs. A comparison of TFs expressed in mouse CR and ETC cells but not in other OB glutamatergic neurons (see Methods) showed that mouse CR and ETC cells expressed different sets of TFs overall, but shared the expression of many TFs usually associated to CR identity (fig. S9D). Of these, *Nr2f2*, *Tp73*, *Dach1*, *Ebf3*, *Zeb2*, *Lhx1*, *Nr2f1*, *Tbr1*, *Id4* and *Hivep1* were expressed also in chicken CR cells and in salamander CR and ETCs (zebrafish ETCs lacked *Nr2f2*, *Tp73*, *Zeb2* and *Hivep1* expression, but we cannot rule out technical limitations) (Fig. 5A). These results indicate that, at least in tetrapods, many “canonical” CR transcription factors are also expressed in ETCs.

**Figure 5.**
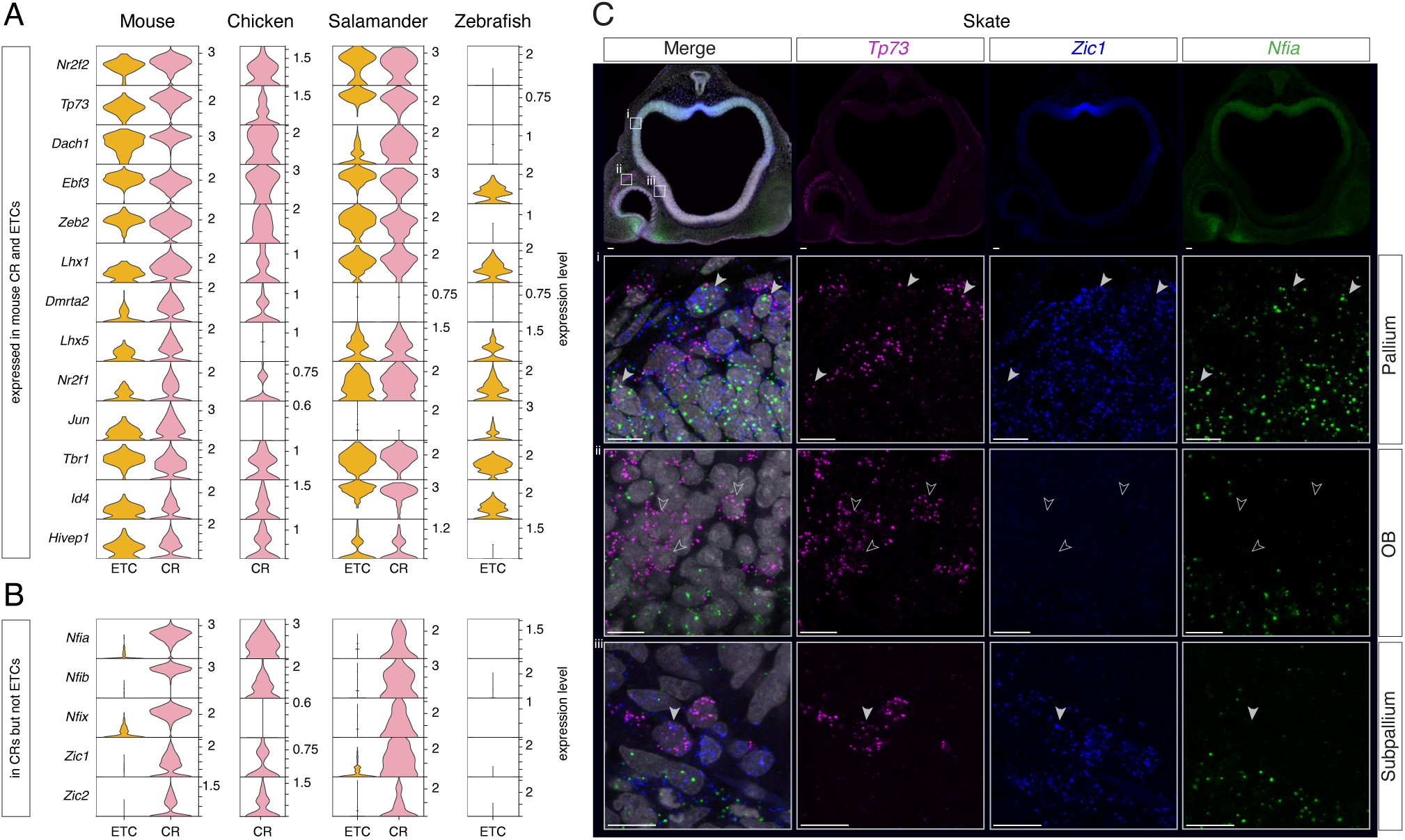
Transcription factor expression in CR cells and ETCs across vertebrates. (A) Top: Violin plots showing expression of TFs expressed in both mouse CR cells and ETC compared to a cell type used as outgroup (OB mitral/tufted cells), and the expression of the same TFs in other vertebrate species. (B) Violin plots showing expression of TFs that differentiate CR cells and ETCs in both mouse and salamander, and the expression of the same TFs in chick CR cells and zebrafish ETCs. Data from ^39,40,44,46,54^ (see Methods for details). (C) HCR triple in situ hybridization for *Tp73*, *Zic1*, and *Nfia* in stage 29 skate telencephalon. (i) Coexpression of all three marker genes in pallial neurons suggesting a CR identity; (ii) *Tp73*+ in OB do not coexpress Zic1 and *Nfia*; (iii) coexpression of *Tp73* and *Zic1* but not *Nfia* in subpallium.

Therefore, we asked whether the molecular identities of ETC and CR cells are defined by the expression of different sets of TFs, *i.e.* if these two cell types are genetically individuated ^55^. Specifically, we looked for TFs that could consistently discriminate between ETC and CR cell identities across vertebrates. First, we identified TFs that had conserved differential expression between ETC and CR cells in both salamander and mouse, and then assessed their expression in chicken CRs and zebrafish ETCs (see Methods). While we couldn’t find any conserved ETC-specific TFs, we observed that *Zic1*, *Zic2*, and NFI-family TFs were expressed in CR but not in ETCs in mouse and salamander, and that these TFs were expressed in chicken CRs too but not in zebrafish ETCs (Fig. 5B). Notably, in mouse NFI-family TFs are specific to hem-derived CR cells ^33^, suggesting that the CR cells sampled in salamander and chicken are mostly hem-derived, in line with our HCR data.

Finally, we checked for the presence of these CR-specific TFs in skate *Tp73+* neurons (Fig. 5C). *Tp73* neurons in the subpallium expressed *Zic1* but not *Nfia*. Moreover, the *Tp73*+ cells in the OB did not express *Nfia* or *Zic1*, consistent with ETC identity. However, *Tp73* showed co-expression with both *Nfia* and *Zic1* in the skate pallium, indicating that these are CR cells. These combinations of markers in the little skate suggest that the major classes of *Tp73*+ neurons existed in the last common ancestor of jawed vertebrates.

Taken together, the high degree of transcriptome and TFs expression similarity of ETC and CR cells suggests that these cell types have a shared evolutionary origin. Across different vertebrate species, CR cells could be consistently distinguished from ETCs for the expression of *Zic* and NFI transcription factors, suggesting that these TFs were recruited in ancestral vertebrates to differentiate the identities of these two cell types.

### Transcriptomic divergence of vertebrate Cajal-Retzius cells

The presence of CR cells in non-mammals challenges the notion that mammalian-specific cortical traits emerged with the appearance of this cell type in evolution. Alternatively, CR cells might be a conserved cell type with diverging characteristics across species.

Since a key effector of mammalian CR function is the secreted protein Reln, we first asked whether the coexpression of *Tp73* and *Reln* is conserved across vertebrate species. In the developing chick brain, we found robust overlap of *Tp73* and *Reln* in the pallium and hem, corroborating scRNA-seq data (Fig. 6A, fig. S6H). However, stainings for *Tp73* and *Reln* in salamander revealed coexpression only in rare *Tp73* pallial neurons, consistent with scRNA-seq (fig. S7H). Furthermore, there was no coexpression of *Tp73* and *Reln* in the pallium of developing skates and zebrafish. This is most striking in the developing skate brain, where *Reln* is clearly absent from the dense layer of *Tp73* neurons at the pallial surface, but is expressed robustly elsewhere, including a separate population of pallial neurons (Fig. 6A, fig. S12). This suggests that strong coexpression of *Tp73* and *Reln* may be an amniote innovation, in line with observations in other species ^28^, and that *Reln* is not a conserved marker of CR cells across all vertebrates.

**Figure 6.**
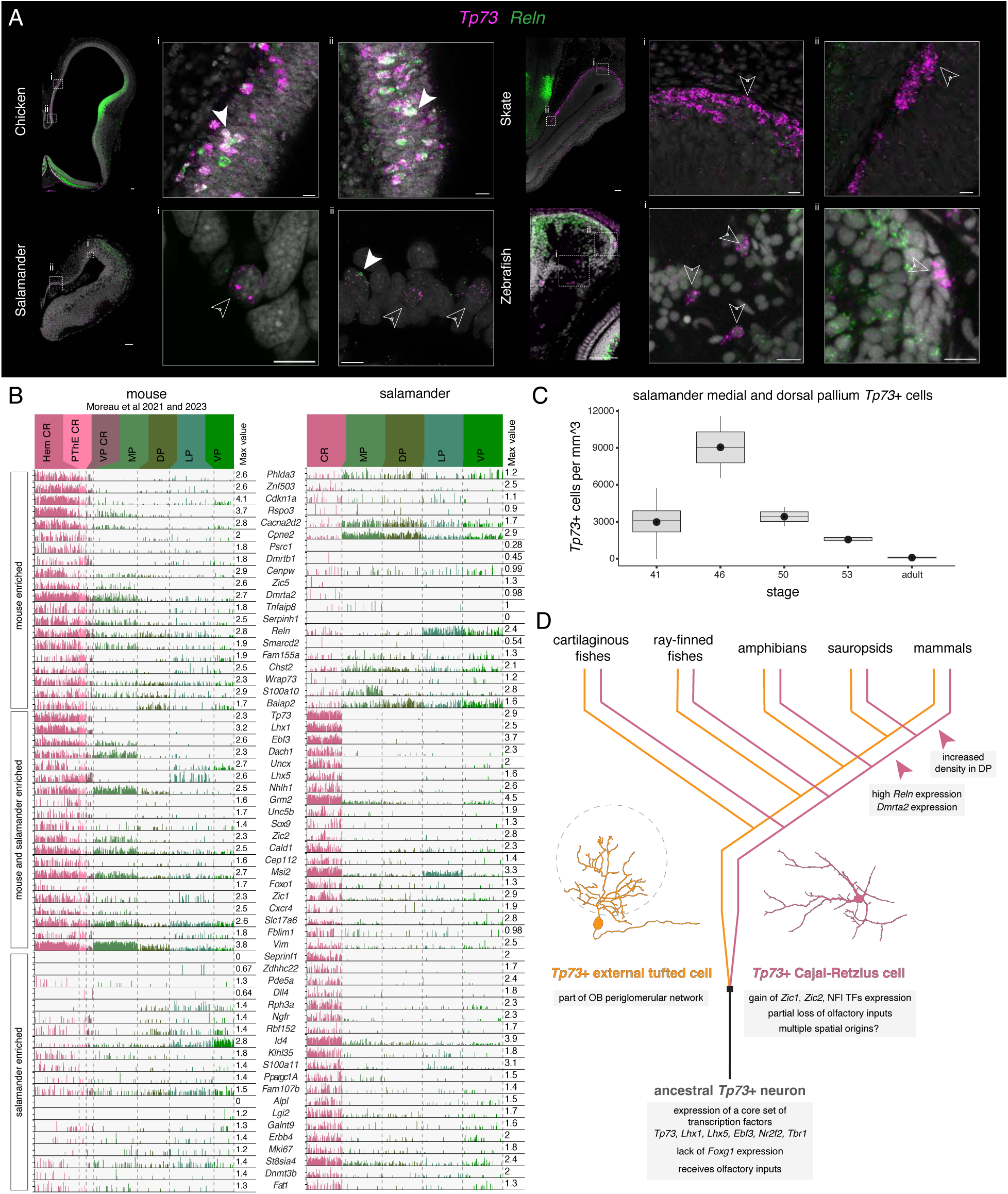
Transcriptomic divergence of vertebrate CR cells. (A) HCRs showing *Reln* (green) and *Tp73* (magenta) expression on coronal sections of developing stage E6 chick, st.50 salamander and 6 dpf zebrafish brains, and an horizontal section of a stage 29 skate brain. Closed arrowheads indicate examples of *Tp73*+ *Reln+* cells. Open arrowheads indicate examples of *Tp73*+ *Reln*-negative cells. Scale bar 50um (Coronal Section) 10 um (Zoom Panel). (B) Expression of marker genes that differentiate CR cells from other pallial glutamatergic neurons (ETCs excluded) in mouse and salamander developmental datasets. The plot shows the top 20 genes enriched in mouse but not salamander CR cells; the top 20 genes enriched in both species, and the top 20 enriched in salamander but not mouse CR cells (see Methods). Mouse data from Moreau et al 2021 and Moreau et al 2023 ^29,33^. (C) Quantification of the density of *Tp73*+ cells in salamander medial and dorsal pallium across developmental stages. Black dot represents average value. n= 3 animals for stages 36-50 and adult, n=2 for stage 53. (D) Proposed evolutionary relationship between CR cells and ETCs. Vertebrate phylogenetic tree with cell type typogenetic tree superimposed (inspired by^74^). Schematics of ETC and CR cells (dorsal view) based on^75,76^. Dotted circle represents OB glomerulus. Abbreviations: DP, dorsal pallium; LP, lateral pallium; MP, medial pallium; VP, ventral pallium

To identify other species-specific specializations, we searched genes with conserved and species-enriched expression in salamander and mouse CR cells. We computed in each species the differentially expressed genes between CR cells and other glutamatergic neurons in the developing pallium (developing mouse data from Moreau et al. ^29,33^); salamander CR cells were defined here as the mature neurons that integrate with mouse CR cells (Fig. 4) and ETCs were excluded (see Methods). As expected, the top shared markers included *Tp73*, *Ebf3, Zic1, Zic2* and other TFs (Fig. 6B). In addition, we found that the glutamate transporter *Slc17a6* (*Vglut2*), the glutamate receptor *Grm2*, and general markers of immature neurons (*Msi1*, *Sox9*, *Vim*) in CR cells from both species. *Cxcr4* and *Zic2*, essential for the proper migration and positioning of CR cells in mice ^56,57^, were also expressed in salamander CR cells, suggesting the evolutionary conservation of molecular mechanisms of CR migration. The Netrin receptor *Unc5b* was also expressed in both salamander and mouse CR cells. Importantly, *Nectin-1* (encoded by *Pvrl1*), *N-cadherin* (*Cdh2*) and *Mllt4* (*Afdn*) were also expressed in CR from both species (fig. S13); in mouse, these adhesion molecules are crucial for the interaction between CR cells and cortical neurons during radial migration ^58^.

Interestingly, salamander and mouse CR cells expressed different sets of synaptic genes, with *Lgi2, Erbb4, Rpha3, Zdhhc22*, and *St8s1a4* enriched in salamander, and *Cacna2d2*, *Baiap2*, *S100a10*, and *Cpne2* enriched in mouse (Fig. 6B). This divergent use of synaptogenesis genes may be due to differences in wiring specificity, genetic drift, relative differences in maturation state^59^, or some other species-specific adaptation. Furthermore, salamander and mouse CR cells differed in the expression of neurotrophic, cell cycle, and apoptotic genes, with *Ngfr*, *Serpinf1*, and *Erbb4* enriched in salamander, and *Cdkna1a* (p21), *Phlda3*, and *Tnfaip8* enriched in mouse (Fig. 6B), suggesting diverging mechanisms for the regulation of neuronal survival. Notably, the transcription factor *Dmrta2* was expressed in mouse but not salamander CR cells (see also Fig. 4G). *Dmrta2* knock-out mice have significantly reduced CR abundance ^60^, suggesting that gain of *Dmrta2* expression in amniote CR cells may have contributed to their higher number and density.

Finally, we reasoned that if CR cells have some role during pallial development also in non-mammals, their density should be highest during neurogenesis. This is clearly the case in the little skate (compare Fig. 1E and fig. S2). After counting *Tp73*+ neurons in the salamander medial and dorsal pallium, which are mostly CR cells, we saw that their density peaks during early pallial neurogenesis ^38,40^ (stage 46, with ∼9000 cells per mm^3), and gradually decreases later (Fig. 6C). Taken together, our data indicate that non-mammalian CR cells may be involved in early pallial development, but use a divergent set of molecular effectors including Reln.

## Discussion

Our molecular comparisons in fish, amphibians, birds, and mammals indicate that *Tp73*+ Cajal-Retzius cells are an ancient cell type in vertebrates. These results place CR cells as part of the broader family of *Tp73*+ neuron types. We identify two conserved classes of *Tp73*+ neurons in vertebrates, with distinct transcriptomes and spatial distribution. A first class (*Tp73*+ *Ebf3*-cells) comprises cells in the PThE and basal forebrain. The PThE is the developmental source of several types of glutamatergic neurons that then migrate to other brain regions, including the preoptic area and hypothalamus ^9,61,62^. We suggest that the *Tp73*+ *Ebf3*-cells we identified in salamander and zebrafish include the PThE and cells that have migrated to the basal forebrain. A second conserved neuronal class is defined by the coexpression of *Tp73* and *Ebf3*; these neurons are located in the olfactory bulb and in the pallium, particularly in the hem region. Further transcriptomic analyses indicate that these cells belong to two distinct types, the *Tp73*+ external tufted cells and the *Tp73*+ Cajal-Retzius cells. Our results, establishing that CR cells existed at least in the last common ancestor of jawed vertebrates, are largely in line with a recent scRNAseq study reporting CR cells in the shark *Scyliorhinus canicula* ^63^.

In mammals, CR cells originate from multiple forebrain regions, and *Tp73* is expressed in all subtypes except the ventral-pallium derived CR cells. Interestingly, all the CR cells we identified in scRNAseq datasets from non-mammalian species express markers of mammalian hem-derived CR cells such as NFI transcription factors. Although the hem might be the most prominent source of CR cells outside mammals, we cannot exclude the existence of septum and PThE-derived CR cells in non-mammals, as suggested by staining data for *Ccno* and *Gmnc* in salamander (fig. S7). The evolution of ventral-pallium derived CR cells, which do not express *Tp73*, remains unresolved. In our cross-species comparisons, only *Tp73*+ neurons had high transcriptomic similarity to mouse CR cells, but it is still possible that *Tp73*-negative CR cells exist in non-mammals but are extremely rare and thus were not sampled.

The non-mammalian CR cells show their highest density in the developing brain, suggesting they may play some role in pallial development like their mammalian counterparts. What exactly this ancestral CR function might be remains a matter of speculation, and will need to be addressed by functional studies. Here we find that *Reln* expression is not a conserved feature of vertebrate CR cells. This suggests that, if CR cells have a conserved function in all vertebrates, this function is Reln-independent. The gain of *Reln* expression late in CR cells evolution may have contributed to the emergence of mammalian-specific cortical features, such as radial migration and inside-out corticogenesis ^1,6^. Indeed, *Reln* deletion in mouse *Tp73*+ CR cells affects cortical lamination, but does not recapitulate the other defects caused by CR cells genetic ablation ^20,64^. Therefore, the regulation of radial migration by CR-derived Reln may have evolved with the transition from outside-in to inside-out corticogenesis, an innovation that took place in mammalian ancestors ^40^.

Besides regulating radial migration, mammalian CR cells have prominent roles in the assembly and maturation of neural circuits in the hippocampus ^15,21–23^ and neocortex ^24^. In neocortical layer 1, CR cells are part of large networks of spontaneously-active neurons ^25^, and regulate the spatial distribution of superficial layer interneurons and dendritic maturation in pyramidal neurons ^8^. In mammals, amphibians, non-avian reptiles, and certain fish species, the superficial neuropil layer (layer 1 in mammals) is a conserved site of integration of long-range inputs from cortical and subcortical areas, including sensory inputs relayed by thalamus ^46,65^. Thus, the modulation of circuit assembly might have been an ancestral function of CR cells ^17,66^. Intriguingly, thalamic spontaneous activity regulates CR density in mouse layer 1 ^17^. It is tempting to speculate that the density of CR cells coevolved with the presence of sensory thalamic projections to the pallium, which are scarce in amphibians and seem more prominent in cartilagineous fishes ^67,68^, where CR cells are particularly abundant during development.

Finally, our results reveal a surprising degree of transcriptomic similarity between CR cells and OB *Tp73*+ ETCs. Several observations suggest that CR cells and ETCs are sister cell types, evolved from an ancestral cell type present in the last common ancestor of vertebrates (or earlier) (Fig. 6D). First, in mouse and salamander these two cell types share the expression of a long list of TFs previously thought to be CR-specific, indicating that their transcriptomic similarity is based on a largely shared regulatory program, therefore is not likely to be a result of convergent evolution. Second, lineage tracing experiments in mice indicate that some ETCs are born from pallial progenitors and then migrate to the OB, suggesting that CR and ETC cells may have a shared developmental origin ^50,51^. Third, CR and ETCs have similar electrophysiological properties: they receive strong inputs from GABAergic interneurons, display synchronous bursting, have prominent inward rectifying currents, and are connected by gap junctions ^4,52,53,69–71^. Fourth, both cell types have strong connections with the olfactory system: ETCs are directly postsynaptic to olfactory sensory neurons, whereas CR cells in the lateral olfactory tract, which function as guidepost neurons during development, seemingly belong to a population of lateral olfactory tract cells that receive second-order olfactory inputs from the bulb ^72^. We thus propose that CR and ETCs may have evolved by duplication and divergence from cells involved in early olfactory processing in vertebrate ancestors. Our TF analysis suggests that CR cells may have evolved by neofunctionalization, by acquiring the expression of *Zic* and NFI transcription factors. This regulatory change may have conferred these cell new signaling functions to regulate circuit assembly in an expanded telencephalon ^73^, further supporting the hypothesis that olfactory circuits in ancestral vertebrates served as a developmental and functional blueprint for the evolving telencephalon (Fig. 6D).

Taken together, our findings also highlight the complex relationships between molecular and phenotypic cell type evolution^55^. Across vast evolutionary time, molecular signatures that define cell identity, such as transcription factor expression, may be preserved even when functional roles have diverged dramatically. Our study demonstrates how in-depth analyses of anatomical, transcriptomic, and developmental data can be used to identify and explain these unexpected evolutionary relationships.

## Acknowledgements

We thank Nandan Nerurkar and Elanur Yilmaz (Columbia University) for providing chick and zebrafish samples, the Columbia ICM team for excellent animal care. We also thank Tosches lab members as well as Oliver Hobert and Hynek Wichterle for feedback throughout the project. Light-sheet imaging was performed with support from the Zuckerman Institute’s Cellular Imaging platform (NIH grant 1S10OD023587-01). Computing resources were provided by Columbia University’s Shared Research Computing Facility (NIH grant 1G20RR030893-01 and NYSTAR contract C090171).

## Author Contributions

M.A.T. supervised the study. E.G., G.G., A.D, M.A.T. analyzed scRNA-seq data, and E.G. performed cross-species comparisons. E.G., G.G., P. B., J A.G., C.K. and J.W. prepared samples for histology. E.G., G.G., and J.W., performed histology and imaging. E.G. and M.A.T. wrote the manuscript, with edits from G.G., J.W., A.D., and C.K.

## Materials and Methods

### Animals

*Salamander*: *Pleurodeles waltl* adults and larvae were obtained from a breeding colony established at Columbia University. Animals were maintained in an aquatics facility at 20°C under a 12L:12D cycle. All experiments were conducted in accordance with the NIH guidelines and with the approval of the Columbia University Institutional Animal Care and Use Committee (IACUC). Breeding was induced naturally by manipulating water level and temperature. Experiments were performed with adult (5-19 months) male and female salamanders, and stage 30, 33, 36, 41, 46, 50, 53, and 55 embryos and larvae, staged according to the Gallien and Durocher 1957 ^77^ staging criteria. Embryos and larvae were anesthetized by immersion in a solution of 0.04% MS-222 (tricaine) for 5 minutes, while for adults a solution of 0.2% MS-222 in tank water was used. For stages 30-46, the brain was extracted directly after anesthesia, while stage 50-55 larvae were first perfused transcardially with 2 ml ice-cold PBS and adults with 10 ml ice-cold PBS and 10 ml of 4% PFA in PBS and decapitated before brain extraction. After overnight incubation in 4% PFA in PBS at 4C, brains were progressively dehydrated to 100% MeOH and stored at −20C.

*Bichir fish*: Adult *Polypterus senegalus* (∼10 cm long) were purchased from commercial suppliers and kept in aquaria at Columbia University at 27C under a 12L:12D light cycle. All experiments were conducted in accordance with the NIH guidelines and with the approval of the Columbia University Institutional Animal Care and Use Committee (IACUC). Individuals were anesthetized using 0.4% MS-222 in tank water and then perfused transcardially with 20 ml ice-cold PBS and 10 ml ice-cold 4% PFA in PBS. After decapitation, the brain was extracted and fixed overnight in 4% PFA in PBS at 4C and dehydrated to 100% MeOH for storage at −20C.

*Chicken*: Fertilized eggs of White Leghorn Chicken (*Gallus gallus*) (Charles River) were incubated in a humidified chamber at 38°C, and embryos were staged according to^78^. Brains were dissected in cold PBS from embryos at E6, E8 and E16 stages. E16 embryos were perfused transcardially with 10ml cold PBS before dissection. Brains were immediately fixed overnight in 4% PFA in PBS at 4C, then dehydrated to 100% MeOH and stored at −20C.

*Zebrafish*: All zebrafish experiments were performed in accordance with the applicable regulations and approved by the Institutional Animal Care and Use Committee (IACUC) at Columbia University (protocol numbers AC-AABN3554 and AC-AACB9700). Wild-type AB zebrafish embryos at 2dpf and 3dpf, larvae at 6dpf and 15dpf, juvenile 45dpf juvenile, and adults of 3mpf and 6mpf were used. 2dpf and 3dpf (treated with PTU for depigmentation) embryos as well as whole 6dpf and 15dpf larvae were fixed overnight in 4% PFA. Additionally, whole brains from 6dpf, 15dpf, 45dpf, 3mpf and 6mpf zebrafish were also dissected after euthanization, and fixed in 4% PFA for overnight at 4°C. After fixation, next day samples were washed several times with PBS, then subjected to dehydration step by series of incubation in 25% MetOH - 75% PBS, 50% MetOH - 50% PBS and 75% MetOH - 25% PBS solutions for 10 minutes each, and finally transferred to 100% MetOH for storage at −20°C.

*Skate*: Little skate (*Leucoraja erinacea*) embryos were obtained from the Marine Resources Center at the Marine Biological Laboratory (MBL) in Woods Hole, MA, USA. All skate experiments were conducted according to protocols approved by the Institutional Animal Care and Use Committee of the MBL. Embryos were reared to stages S27, S29, S32 and hatchling stage as described in Gillis et al ^79^. Embryos were euthanized with an overdose of MS-222 (1 g/L in seawater), fixed overnight in 4% PFA in PBS and stored in 100% MeOH at −20C. Hatchlings were anesthetized by immersion in 0.5% MS-222 for 30 min, then perfused transcardially with 20 ml cold PBS and 10 ml of 4% PFA in PBS. After decapitation, and brain extraction, brain tissue was fixed overnight in 4% PFA in PBS at 4C and dehydrated to 100% MeOH for storage at −20C.

### Analysis of scRNA-seq datasets

Selection of mature neurons from the developmental salamander dataset.

To analyze salamander neurons and compare them with neurons from other species, we subsetted the full developmental dataset to include only mature neurons. This selection allowed us to include data spanning multiple developmental stages where we expected CR cells to be present, while preventing neuronal maturation from being the dominant feature driving gene selection and downstream analyses.

Mature neurons from the full larval salamander data set (Deryckere et al 2025) ^40^ were identified using the AddModuleScore_UCell function within the UCell package ^80^, and using the following signature: positive genes: *Snap25, Syp, Stx1a, Mapt, Eno2, Map2*, and negative genes: *Pcna* and *Mki67*. Cells were annotated as mature neurons if they had a module score above 0.4. Mature neurons were only kept from clusters that contained at least ⅔ mature neurons. This was done to remove sparse cells that had a positive neuronal signature but were in primarily non-neuronal clusters. In order to ensure all mature *Tp73*+ neurons were present for integration, special care was given to these cells when identifying mature neurons. For any cell that was in the *Tp73* subsetted object (see below), only cells in cluster clusters im1 and im2 were excluded from the full mature neuron object.

Clusters were annotated by brain region (telencephalon, diencephalon, and midbrain) based on marker genes curated from literature. Clusters were categorized as Glutamatergic, Gabaergic or Other based on the expression of *Slc17a6*, *Slc17a7*, *Gad2* and *Th*.

Selection of mature neurons from the developmental mouse dataset.

All cells published in the developing mouse brain atlas (La Manno 2021) ^41^ were filtered to remove cells with less than 2000 UMIs and with a percentage of mitochondrial genes above 10%. Cells annotated as “Hindbrain” were then removed because hindbrain cells were not included in the salamander and zebrafish datasets. Cells annotated by La Manno et al. as “Bad cells” were also removed. The remaining dataset was normalized using the Seurat SCTransform function while regressing for percent.mt, nCount_RNA, Age, and CellCycle. Clusters were computed using the Seurat FindClusters function with the parameters dims=35 and res = 8. Individual neurons were determined to be mature based on receiving a class annotation as “Neuron” from Lamanno et al. Clusters that contained at least ⅔ mature neurons were kept for further analysis. Regional identity of a neuronal cluster was established using marker genes curated from literature. Clusters were categorized as Glutamatergic, Gabaergic or Other based on the expression of *Slc17a6*, *Slc17a7*, *Gad2* and *Th*.

Selection of mature neurons from the developmental zebrafish dataset.

The published scRNA-seq atlas from the developing zebrafish brain by Pandley et al was filtered to keep mature neurons according to their annotated clusters. First, all cells sampled from adult stages were removed, leaving 6 and 15dpf samples. Olfactory bulb neurons were kept as OB. Subpallium_01 to Subpallium_08 clusters were re-annotated as Telen GABA. Pallium_01 to Pallium_08 clusters were re-annotated as Telen Glut. Pallium_09 was re-annotated as Diencephalon based on the expression of *lhx1a*, *tbr1b*, *barhl2*, and lack of *tac3b* expression, which uniquely correlates with PThE based on results in MapZeBrain.org ^81^. Preoptic area (PoA) and Habenula clusters were also annotated as Diencephalon. These subsetted neurons were normalized using the Seurat SCTransform function while regressing for stage. The top 3000 variable genes were used for downstream analysis, and the first 35 PCs were used for UMAP embedding.

#### Salamander *Tp73*+ neurons dataset

The Salamander *Tp73*+ neurons dataset includes cells selected from two datasets:

1. From the developmental salamander brain dataset (Deryckere et al)^40^: clusters were computed from the full salamander developing brain object using the FindClusters function in Seurat with the res parameter set to 4. Clusters where *Tp73* was expressed in more than 5% of cells, and where the average scaled expression of *Tp73* was above 0.1 were selected for further analysis.
2. From the adult salamander forebrain dataset (Woych 2022)^46^: two *Tp73*+ clusters (TEGLU31 and DIMEGLU6) were selected after the application of the same filtering criteria described above.

These selected developmental and adult *Tp73*+ clusters were then merged, reclustered and further filtered to keep only *Tp73*+ neurons, by removing choroid plexus, epithelial cells, progenitors, and some *Tp73*-negative neurons that made it through the previous round of filtering (details of filtering in code, and fig. S6). These final subsetted 982 *Tp73*+ neurons were normalized using the Seurat SCTransform function. The top 1000 variable genes that were expressed in at least 2.5% of cells were used for downstream analysis, and the first 20 PCs were used for clustering and UMAP embedding, with clusters calculated using a resolution of 1.55.

#### Mouse *Trp73*+ neurons dataset

*Trp73*+ forebrain neurons were identified from the full Allen Brain Cell atlas (Yao 2023)^39^ based on the expression of *Trp73* and their annotation as neurons (*Trp73* is also expressed in some non-neuronal cell types). Groups were selected regardless of their level in the taxonomy. The final selection included: subclass 118 ADP-MPO Trp73 and cluster 2111 LHA Barhl2 Glut_2 (grouped as PV), supertype 0132 OB Eomes Ms4a15 Glut_3 (external tufted cells), and subclass HPF CR Glut (labeled as CR). Raw count matrices and metadata were downloaded from the Allen Brain Cell atlas portal in March 2024. These data are organized by dissected regions. Therefore each raw data matrix was filtered to keep only cells belonging to the selected *Tp73*+ groups, and cells sequenced with the 10x v3 chemistry. This produced a final dataset of 2839 cells. This dataset was then normalized using the Seurat NormalizeData function, and the top 3000 variable genes were computed for downstream analysis.

Calculation of Gene Specificity Index (GSI) correlations.

GSI correlations between cell classes defined by region and primary neurotransmitter (see above) were computed using the method established in Tosches et al 2018. For the salamander-mouse comparison, differentially expressed genes were identified after running the Seurat FindAllMarkers function, using a 0.75 logfc.threshold and an adjusted p-value of 0.01. One-to-one orthologs were identified as described in Woych et al 2022 ^46^. For the zebrafish-salamander comparison, differentially expressed genes were identified based on a 1 logfc.threshold and adjusted p-value of .001.

Because teleost ancestors underwent a whole-genome duplication and zebrafish have several recently duplicated paralogs of tetrapod genes, an alternative to the standard one-to-one orthologue filtering was implemented. Eggnog ^82^ was run using taxonomic scope = vertebrates, Orthology restrictions = Any. For zebrafish genes that were orthologous to multiple different vertebrate gene names, only the one with the lowest e-value was kept. For multiple zebrafish genes that mapped to the same vertebrate gene name, the counts were summed and the vertebrate gene name was assigned for cross species comparison.

#### Marker gene enrichment analysis

To determine CR cell marker genes, CR cells were compared to other mature neurons (see above) in the La Manno et al. ^41^. This list was filtered to keep genes that had an avg_log2FC >2 and a difference in percent of cells expressing the gene greater than .2 between the CR cells and other neurons. This list was then brought into salamander gene naming convention as described above. Orthologous genes were kept for module score calculation in salamander if they showed expression in the Seurat object (*i.e.* were listed as a feature). This final list of 49 marker genes was used as input to Seurat’s AddModuleScore function. For the zebrafish analysis, the top marker genes for salamander *Tp73*+ neurons were computed using the same approach, yielding 42 marker genes.

#### Cross-species scRNA-seq integration

For cross species integration, genes between species were brought into the same naming convention as described above. Mature neurons from developing salamander, mouse and zebrafish were identified as described above, and further subsetted to include only forebrain neurons in the case of salamander and mouse. For Seurat CCA integration of mature neurons, we used functions within Seurat as follows: We selected 2000 integration features using “SelectIntegrationFeatures”, and then subsetted this list for genes that were variable (based on variable.features.n = 4000) and expressed in at least 0.25% of cells in both species separately. The resulting list of integration features was passed to PrepSCTIntegration(). Then pairs of mutual nearest neighbors (anchors) were identified using the FindIntegrationAnchors function with the arguments reduction=“cca” (Canonical Correlation Analysis) and normalization.method = SCT, before ultimately performing integrations using IntegrateData with 90 dims (in the case case of mouse to salamander) and 80 dims (in the case of salamander and zebrafish). The same dimensions used for integration were used for UMAP embedding. Clusters were calculated for the mouse-salamander integration using a clustering resolution of 1. Clusters were calculated for the salamander-zebrafish integration using a resolution of 2.

The same pipeline was used for integration of *Tp73*+ neurons with the following exceptions: Initial identification of variable genes used a threshold of 8000 for each species, that needed to be expressed in 1% of cells within each original species (for zebrafish mouse integration, at least .1% of zebrafish cells to account for difference in sequencing depth), which was used to filter a list of 5000 integration features. 50 dimensions were used for integration and UMAP embedding, and clustering was performed using a 0.1 clustering resolution for chicken and salamander with mouse, and .05 for zebrafish with mouse.

As an alternative to co-clustering analysis, we analyzed cell type neighborhoods in the shared nearest-neighbor (SNN) graph computed by Seurat after integration. The SNN graph is the output of the FindNeighbors function and the input used for clustering. For each species and cell type, we counted the number of neighbors from the other species in the integrated SNN graph, and classified them according to cell type identity. This produced two matrices (*i.e.* salamander neighbors of mouse cells, and mouse neighbors of salamander cells) that were normalized by the total number of neighbors per source cell type, and then averaged to obtain the SNN proximity score.

#### Identification of differentially expressed genes between mouse *Tp73+* neuron types

Identification of marker genes for mouse *Tp73+* CR and ETC cells were determined relative to *Tp73*-negative mitral and tufted cells (MT), which were the remaining supertypes in the 03-OB-CR Glut cell class in the Allen Brain Cell atlas. These MT supertypes were added to the mouse *Tp73*+ neurons object (described above), to generate a new object with 3516 cells, and were normalized using the Seurat SCTransform function. The top 3000 variable genes were used for downstream analysis. Marker genes were identified using the FindMarkers function within Seurat. Genes were listed as upregulated in either CR or ETC if relative to MT cells, they: had avg_log2FC>.5, had over a difference over 25% in the percentage of cells expressing the gene (*i.e.* 36% in CR, 10% in MT), had at least twice the percentage expressing the gene (*i.e.* 81% in CR, 40% in MT), and the differential expression of the gene received a p-value below 0.001. Genes were considered shared between CR and ETC if they were upregulated (relative to MT) in both lists. Genes were considered specific for CR or ETC if they were upregulated for one cell type but not the other, and were lowly expressed in the other cell type (below 25% of cells expressing the gene). Genes were filtered for display in Violin plots based on GO term. In order to show the conservation of gene expression across species, transcription factors that were expressed in both mouse CR and ETC had their expression checked in chick ^54^, salamander ^40,46^ and zebrafish ^44^. TFs were included in Figure 4G if they were expressed in at least 2 species.

For genes that differentially label CR cells from ETCs, Seurat FindMarkers was run to compare CR cells against ETCs in both the salamander and mouse data with test.use=“wilcox”. Transcription factors that had over 25% difference in percent expression, an Avglog2FC above 2, below 35% expression in ETCs, and an adjusted p.value below .05 were considered CR specific markers for a species. The intersection of CR specific TFs between salamander and mouse is shown in figure 4. The same analysis was run to identify ETC specific markers across species but no transcription factors met the above significance threshold in both salamander and mouse.

#### Identification of differentially expressed CR markers between mouse and salamander

Comparisons of mouse and salamander CR marker genes were performed after computing differentially expressed genes between CR cells and other glutamatergic pallial neurons in developmental datasets. We filtered our data to ensure that the only *Tp73*+ cells were CR cells because we did not want to identify markers that are unique to basal forebrain or ETC *Tp73+* neurons. In salamander, the *Tp73*+ neuron object was filtered to remove adult cells, as well as cells in clusters im1 and im2. Then, based on the *Tp73* integration described above, cells that clustered with mouse CR cells were labeled as salamander CR. This included all of m1.1 as well as a portion of m1.2. These 87 salamander CR cells were merged with 100 randomly selected mature neurons from each of MP, DP, LP and VP to create an object with 487 total cells. To obtain high quality developmental CR cells in mice we relied on published data from Moreau et al 2021 ^33^ for CR cells originating from hem, PThE, septum and ventral pallium, as well as Moreau et al 2023 ^29^ for hem-derived CR cells. All cells labeled as CR in Moreau et al 2021 were taken for further analysis as well as all clusters that could be annotated as dorsal, lateral or ventral pallium based on marker genes provided in their results. Data from Moreau 2023 were subsetted to only include CR cells and Pallial neurons, which in this case based on dissection and marker genes were all medial pallium. These cells were normalized using the SCTransform function while regressing for RNA count. The top 3000 variable genes were used for downstream analysis, and the first 50 PCs were used for clustering analysis and UMAP embedding; Louvain clustering was performed with a clustering resolution of 2. A random sample of 100 cells each was taken from the most mature cluster of Cell.state Cajal_retzius_neurons_LN and Pallial_neurons_LN based on the value of LN_signature1. Combining these cells with the subset from Moreau et al 2021 resulted in an object with 449 cells.

We then found differentially expressed genes between *Tp73*+ CR cells and Pallial glutamatergic neurons using the FindMarkers function with min.pct = .0001, logfc.threshold= .001, and test.use=”wilcox”. We labeled a gene as a CR marker if it had an average log-fold-change greater than 1 in CR cells relative to pallial neurons. We also filtered by adjusted p-value, only accepting DE genes with an adjusted p-value below 0.000001. Genes were categorized as species enriched CR genes if they appeared in the list of CR upregulated genes for their species but not in the list for the other, and the other species had below 33% expression of that gene in their CR cells. Genes were categorized as shared if they were in the list of upregulated CR genes for both species. For the plot in Figure 5A, the top 20 genes of each category were selected based on order of average log-fold change relative to the other pallial neurons. It is important to note that due to the selection of pallial glutamatergic neurons as the term of comparison, some known CR marker genes (such as the NFI TFs) were absent from the list, due to their widespread expression in medial and dorsal pallium.

### Hybridization chain reaction (HCR) in situ hybridizations

#### Whole-mount HCR

Whole mount HCR (hybridization chain reaction^83^ in situ hybridizations were conducted as previously described in ^46,84^. Briefly, whole brains of salamander embryos, larvae and adults, chicken E6 and E8 embryos, zebrafish 15-day larvae, and *Polypterus* adults, as well as whole skate S27 and S29 embryos were delipidated with overnight incubation in 100% Dichloromethane (DCM, Sigma-Aldrich 34856) and bleached overnight with 5% hydrogen peroxide in MeOH. Chicken, *Polypterus* and skate specimens were further incubated in bleaching solution (5% Deionized formamide, 1.5% H2O2, 0.2× SSC in nuclease-free water) for 40 mins under strong light. HCR-3.0 probe pairs were designed using **insitu_probe_generator** ^85^ and synthesized by Integrated DNA Technologies (IDT). Probes were added to Hybridization Buffer (Molecular Instruments) to a concentration of 12nM and incubated for three days at 37C. Samples were washed in Wash Buffer (Molecular Instruments) and SSCT (5xSSC + 0.1% TritonX) before incubation with 60pM snap-cooled HCR hairpins (Molecular Instruments) in Amplification Buffer (Molecular Instruments) for 3 days at 4C. Nuclear staining (DAPI or BOBO-1) was also added to the solution with a 1:1000 dilution. After washes in SSCT and 500 mM Tris-HCl (pH 7.0), samples were embedded in 4% agarose in 500 mM Tris-HCl (pH 7.0) and dehydrated to 100% MeOH overnight, then incubated with 66% DCM - 33% MeOH for 3 hours, 100% DCM for 30 min and 100% Dibenzyl ether (DBE, Sigma-Aldrich 108014) overnight. Cleared whole-brain samples were imaged using a LaVision UltraMicroscope II light-sheet microscope.

After staining and agarose embedding, some whole-mount brains and all skate embryos were embedded in 4% agarose prepared in 500 mM Tris-HCl (pH 7.0), and then sectioned at 70 µm using a vibratome. Sections were incubated in DAPI diluted in Tris-HCl for 30 minutes, then mounted in either Fluoromount-G® (SouthernBiotech) or DAKO fluorescent mounting medium (Agilent Technologies) and imaged with a Zeiss LSM800 laser scanning confocal microscope.

#### HCR on vibratome sections

Brains of E16 chicken embryos and of skate hatchlings, and whole-mount 6-day old zebrafish larvae and S32 skate embryos were processed for HCR on sections. Brains stored in 100% MeOH were gradually rehydrated to PBS, then embedded in 4% agarose in PBS and sectioned using a Leica VT1200S vibratome. 100 µm thick sections of skate hatchling brain and 150 µm thick sections of chick E16 brains were then processed for HCR following^86^. Sections were first treated with bleaching solution (5% Deionized formamide, 1.5% H2O2, 0.2× SSC in nuclease-free water) for 30min under strong light, then rinsed in PBS and permeabilized for 1h with a solution of 1% Triton-X and 1% DMSO in PBS. After rinsing in SSCT sections were pre-hybridized in Hybridization Buffer (Molecular Instruments) for 1.5h at 37C and then incubated overnight with probes at 12nM concentration in Hybridization Buffer at 37C. The following day, samples were washed in Wash Buffer (Molecular Instruments) and then in SSCT, then incubated in Amplification Buffer (Molecular Instruments) for 30 min. Sections were then incubated overnight in the dark at RT with snap-cooled HCR hairpins (Molecular Instruments) at a concentration of 0.3µM in Amplification Buffer. Samples were rinsed several times with SSCT, mounted on glass slides with Fluoromount-G (Invitrogen, 00-4958-02) and imaged with a Zeiss LSM800 laser scanning confocal microscope.

#### Quantification of medial and dorsal pallium *Tp73*+ neurons during salamander development

Coronal sections of pallium across developmental stages were stained and imaged as described above, using the 546 channel for *Reln* and the 647 channel for *Tp73*. Dense *Reln* expression in lateral pallium was used to delineate the boundary between dorsal pallium and lateral pallium. In posterior sections of the telencephalon the ventral boundary of the medial pallium was recognized from the clear separation of the pallium from septum. In more anterior sections, this boundary was recognized from a discontinuity of cell density between medial pallium and septum. For each coronal section, medial and dorsal pallium *Tp73*+ cells were manually counted across the Z width (with a z step of 3 um), and a *Tp73* density was calculated per section. These density data were then averaged for each brain to generate the plot in 5B.

