## Supplementary Figures for "Evolutionary origins and transcriptomic innovations of vertebrate Cajal-Retzius cells"

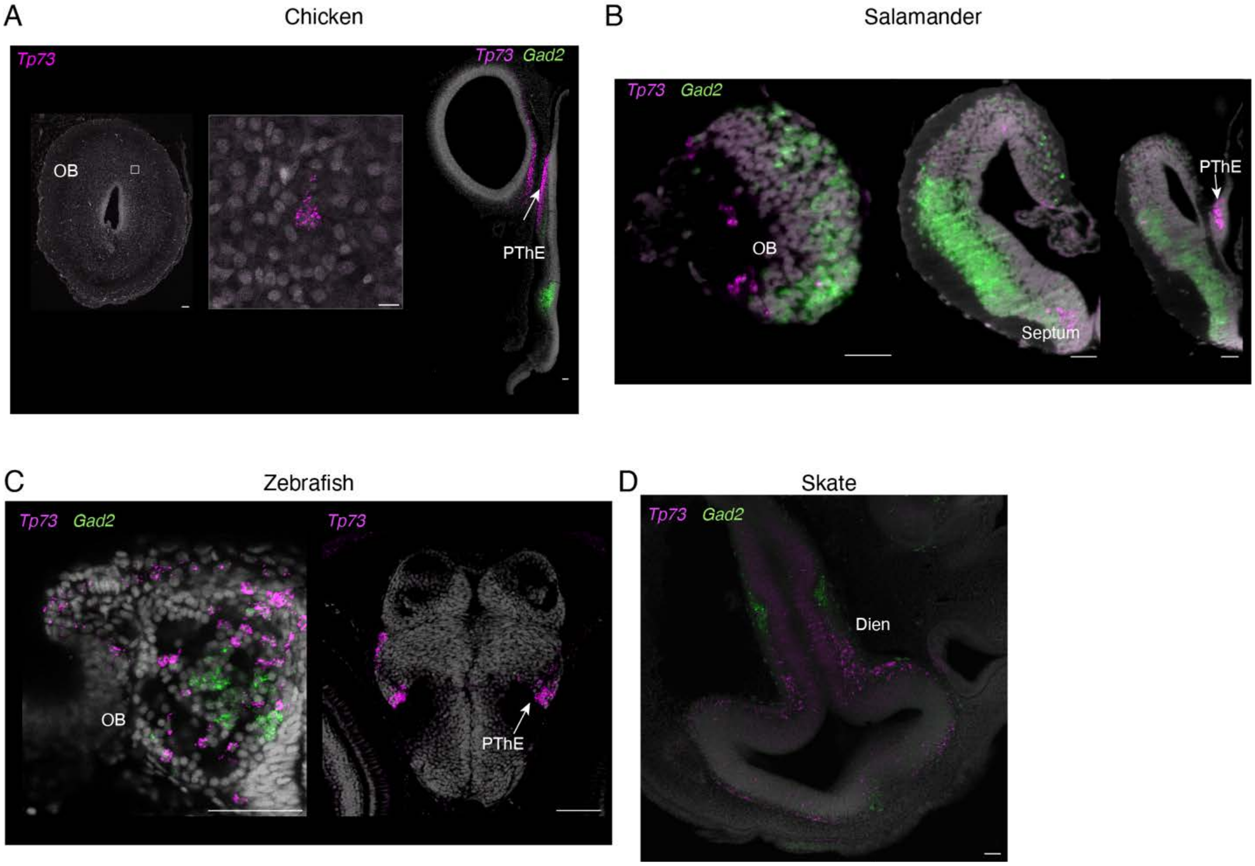


**Supplementary Figure 1. Additional characterization of *Tp73* expression in non-mammalian vertebrates.**
(A) *Tp73* expression in E6 chick OB (left) and PThE (right). *Gad2* labels GABAergic neurons; absence of *Gad2* expression in anterodorsal diencephalon confirms that the *Tp73*+ diencephalic area is the PThE, which is glutamatergic. (B) Optical coronal section of whole mount *Tp73* (magenta) and *Gad2* (green) HCRs in stage 46 salamander larvae, showing *Tp73* expression in the glomerular layer of the OB, in the septum and PThE. (C) Coronal section of 15dpf zebrafish brain showing *tp73* neurons in OB (right) PthE (left; identified by anatomical structure and comparison with Kunst 2019 [87,88]. (D) Horizontal section of stage 27 skate brain, showing *Tp73* expression in *Gad2*-negative cells in the anterodorsal diencephalon, putative position of the PThE. Scale bars 50um.

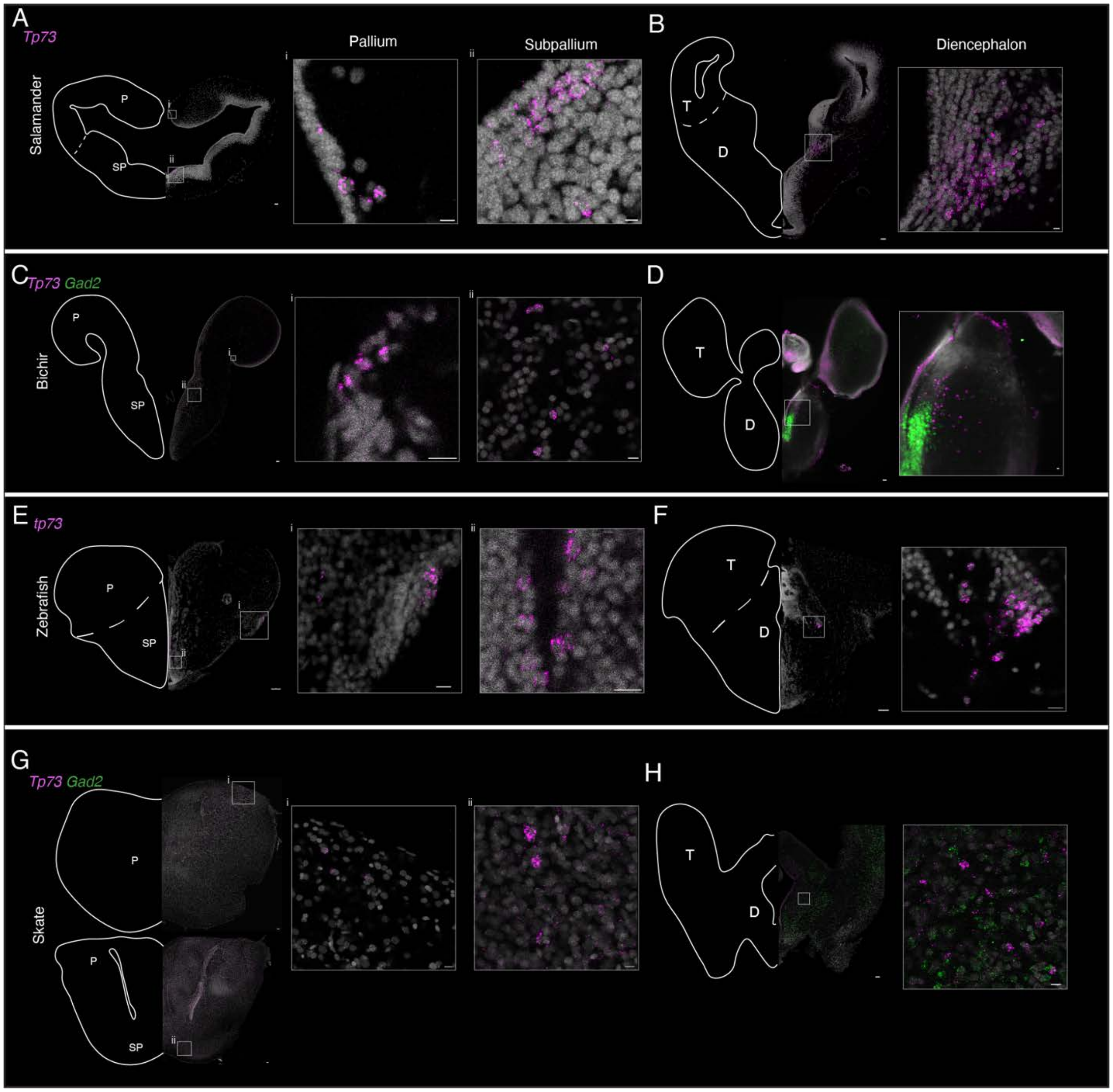


**Supplementary Figure 2. Conserved spatial distribution of *Tp73*+ neurons in adult vertebrate brains.**
HCR *in situ* hybridizations for *Tp73* (magenta) and *Gad2* (green). (A) Coronal section of adult salamander telencephalon showing *Tp73*+ neurons in pallium (hem) and subpallium. (B) Coronal section of adult salamander brain showing *Tp73* expression in the PThE region of diencephalon. (C) Coronal section of bichir telencephalon, showing *Tp73*+ neurons in pallium (hem) and subpallium. (D) Optical coronal section of the bichir brain, showing *Tp73*+ neurons in the putative PThE, identified as a *Gad2*-negative region of the anterior dorsal diencephalon. (E) Coronal section of adult zebrafish telencephalon, with *Tp73*+ neurons in pallium and subpallium. (F) Coronal section of adult zebrafish brain with *Tp73*+ neurons present in thalamic eminence region of diencephalon based on Kenney et al 2021 [87,89]. (G) Coronal sections of adult skate telencephalon at two different rostrocaudal levels, showing scattered *Tp73*+ neurons in

pallium and subpallium. (H) Coronal section of adult skate brain, with *Tp73*+ cells present in putative PThE region of diencephalon. Scale bars 50um for full sections,10 um for zoomed images.

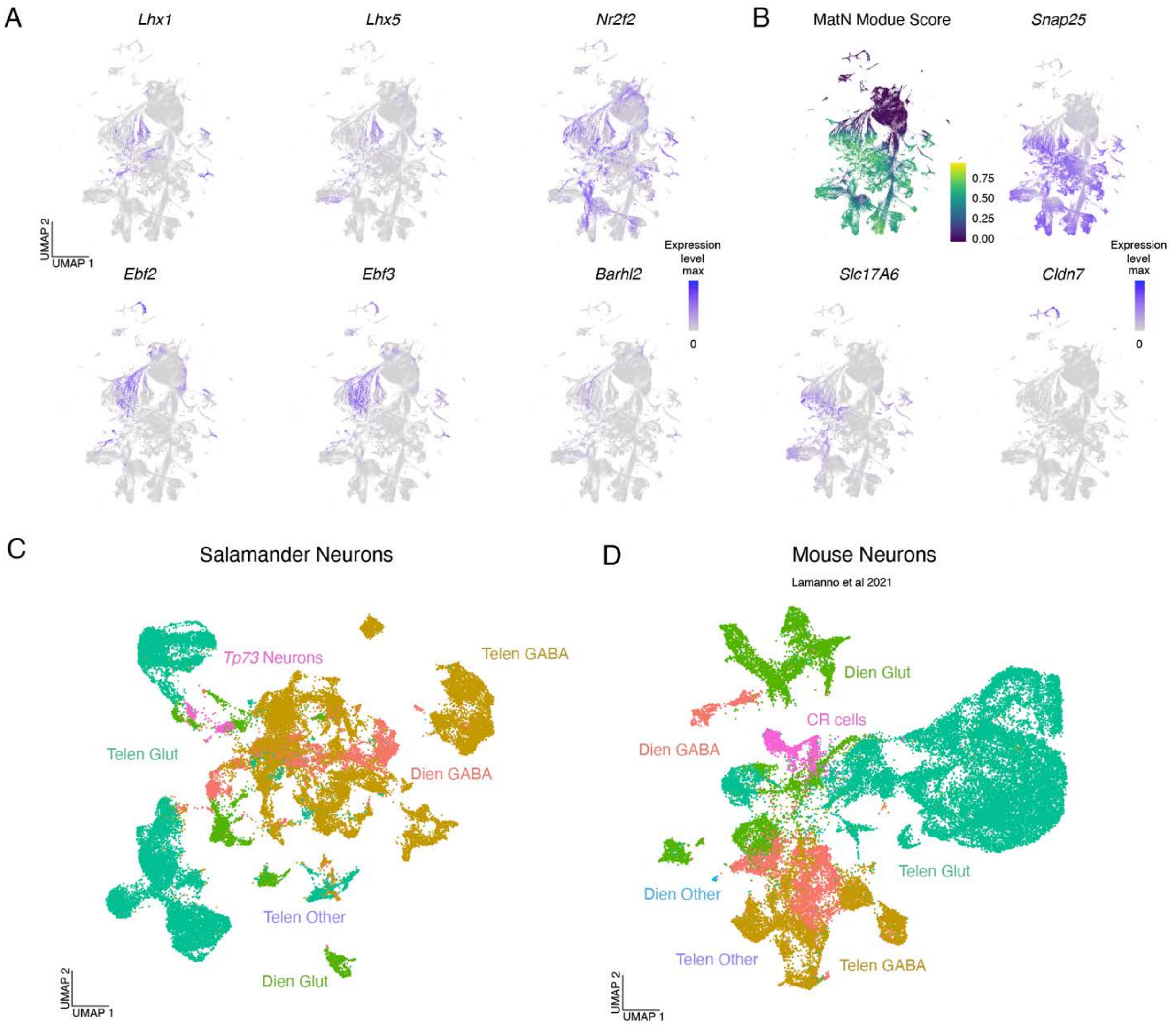


**Supplementary Figure 3. Identification of mature neurons expressing CR marker genes in salamander and mouse scRNAseq datasets.**
(A) Full UMAP showing expression of CR marker transcription factors highlighted in Figure 2B. (B) Top left: UMAP plot of mature neuron UCell Module score used to identify neuronal clusters for subsequent analysis. Module score based on positive expression of *Snap25, Syp, Stx1a, Mapt, Eno2, Map2,* and low expression of *Pcna* and *Mki67*. Other panels: UMAPs showing expression of *Snap25* and *Slc17A6*, confirming that cells highlighted in A are glutamatergic neurons. *Cldn7* expression demonstrates that there is a separate non-neuronal *Tp73* cluster that likely includes the choroid plexus. (C)-(D) UMAPs showing mature neurons classes from the developmental salamander (C) and mouse (D) data (mouse neurons from La Manno et al 2021 [41]). The annotations, based on brain region and primary neurotransmitter, are used for the comparative analyses in Figure 2.

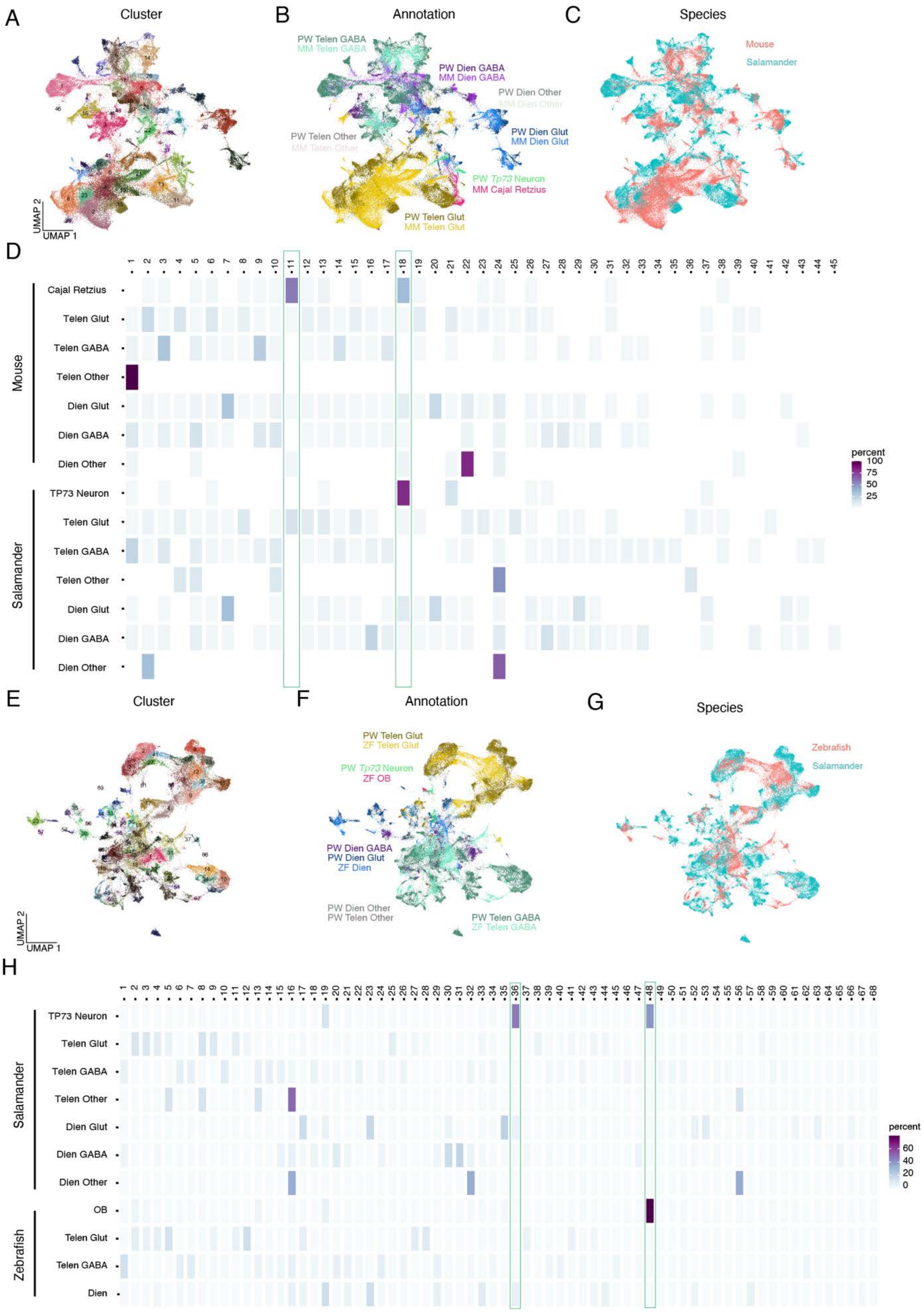
A
Cluster
B
Annotation
C
Species
PW Telen GABA
MM Telen GABA
PW Dien GABA
MM Dien GABA
PW Dien Other
MM Dien Other
PW Dien Glut
MM Dien Glut
PW Telen Other
MM Telen Other
PW Tp73 Neuron
MM Cajal Retzius
PW Telen Glut
MM Telen Glut
Mouse
Salamander
UMAP 1
UMAP 2
D
Mouse
Cajal Retzius
Telen Glut
Telen GABA
Telen Other
Dien Glut
Dien GABA
Dien Other
Salamander
TP73 Neuron
Telen Glut
Telen GABA
Telen Other
Dien Glut
Dien GABA
Dien Other
percent
100
75
50
25
E
Cluster
F
Annotation
G
Species
PW Telen Glut
ZF Telen Glut
PW Tp73 Neuron
ZF OB
PW Dien GABA
PW Dien Glut
ZF Dien
PW Dien Other
PW Telen Other
PW Telen GABA
ZF Telen GABA
Zebrafish
Salamander
H
Salamander
TP73 Neuron
Telen Glut
Telen GABA
Telen Other
Dien Glut
Dien GABA
Dien Other
Zebrafish
OB
Telen Glut
Telen GABA
Dien
percent
60
40
20
0

**Supplementary Figure 4. Integration of salamander scRNA-seq data with mouse and zebrafish data.**
(A-C) UMAPs of Seurat CCA integration of mature mouse neurons from La Manno et al 2021[41] and salamander neurons from Deryckere et al 2025 [40], see also Fig. 2. Cells colored by integrated cluster (A), original annotation (B) and species (C). (D) Matrix describing the composition of each integrated cluster in the salamander-mouse integration. Integrated clusters (11 and 18) with a high proportion of mouse CR cells are highlighted. Integrated cluster 18 also contains the majority of salamander *Tp73*+ neurons. (E-G) UMAPs of Seurat CCA integration of mature zebrafish neurons from Pandey et al 2023 [44] and salamander neurons from Deryckere et al 2025 [40], see also Fig. 2. Cells colored by integrated cluster (E), original annotation (F) and species (G). (H) Matrix describing the composition of each integrated cluster in the zebrafish-salamander integration. Integrated clusters (36 and 48) containing a high proportion of salamander *Tp73*+ neurons are highlighted. Integrated cluster 48 also contains the majority of zebrafish neurons annotated as OB neurons.

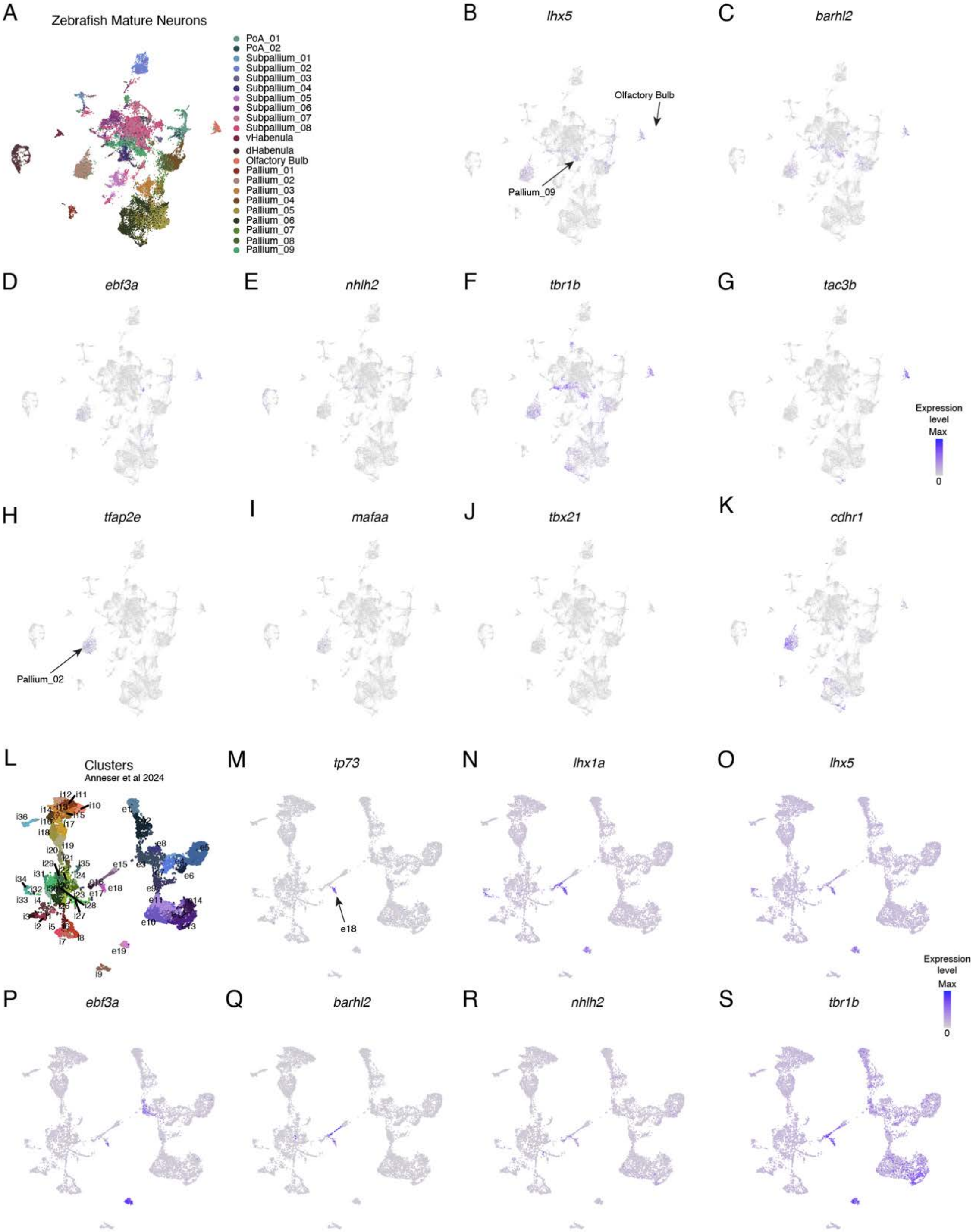

A
Zebrafish Mature Neurons
PoA_01
PoA_02
Subpallium_01
Subpallium_02
Subpallium_03
Subpallium_04
Subpallium_05
Subpallium_06
Subpallium_07
Subpallium_08
vHabenula
dHabenula
Olfactory Bulb
Pallium_01
Pallium_02
Pallium_03
Pallium_04
Pallium_05
Pallium_06
Pallium_07
Pallium_08
Pallium_09
B
lhx5
Olfactory Bulb
Pallium_09
C
barhl2
D
ebf3a
E
nhlh2
F
tbr1b
G
tac3b
Expression level
Max
0
H
tfap2e
Pallium_02
I
mafaa
J
tbx21
K
cdhr1
L
Clusters
Anneser et al 2024
M
tp73
e18
N
lhx1a
O
lhx5
Expression level
Max
0
P
ebf3a
Q
barhl2
R
nhlh2
S
tbr1b

**Supplementary Figure 5. Expression of additional marker genes in zebrafish neurons.**

(A) UMAP for the Pandey et al 2023 [44] dataset showing neurons from 6 and 15 dpf zebrafish. Annotations according to Pandey et al 2023 [44]. (B-G) UMAPs showing expression of key marker genes shown in Fig. 2. *lhx5*, *barhl2*, *nhlh2* and *tbr1b* are expressed in clusters annotated as "olfactory bulb" and "Pallium_09" and are known CR markers. *ebf3a* (a known CR TF) and *tac3b* (specific to zebrafish OB) are expressed in the OB cluster but not in Pallium_09. See Methods for re-annotation of Pallium_09 as diencephalon for integration. (H-K) expression of known marker genes of OB glutamatergic neurons (mitral and tufted cells): tfap2e, mafaa, tbx21(sparsely detected) and cdhr1, that support Pallium_02 being a cluster of glutamatergic neurons in OB. (L) Top left: UMAP showing adult zebrafish neuron clusters from Anneser et al 2024 [45]. (P-S) coexpression of known CR marker genes in cluster e18, along with *tp73*, which is poorly detected in the Pandey et al 2023 dataset [44].

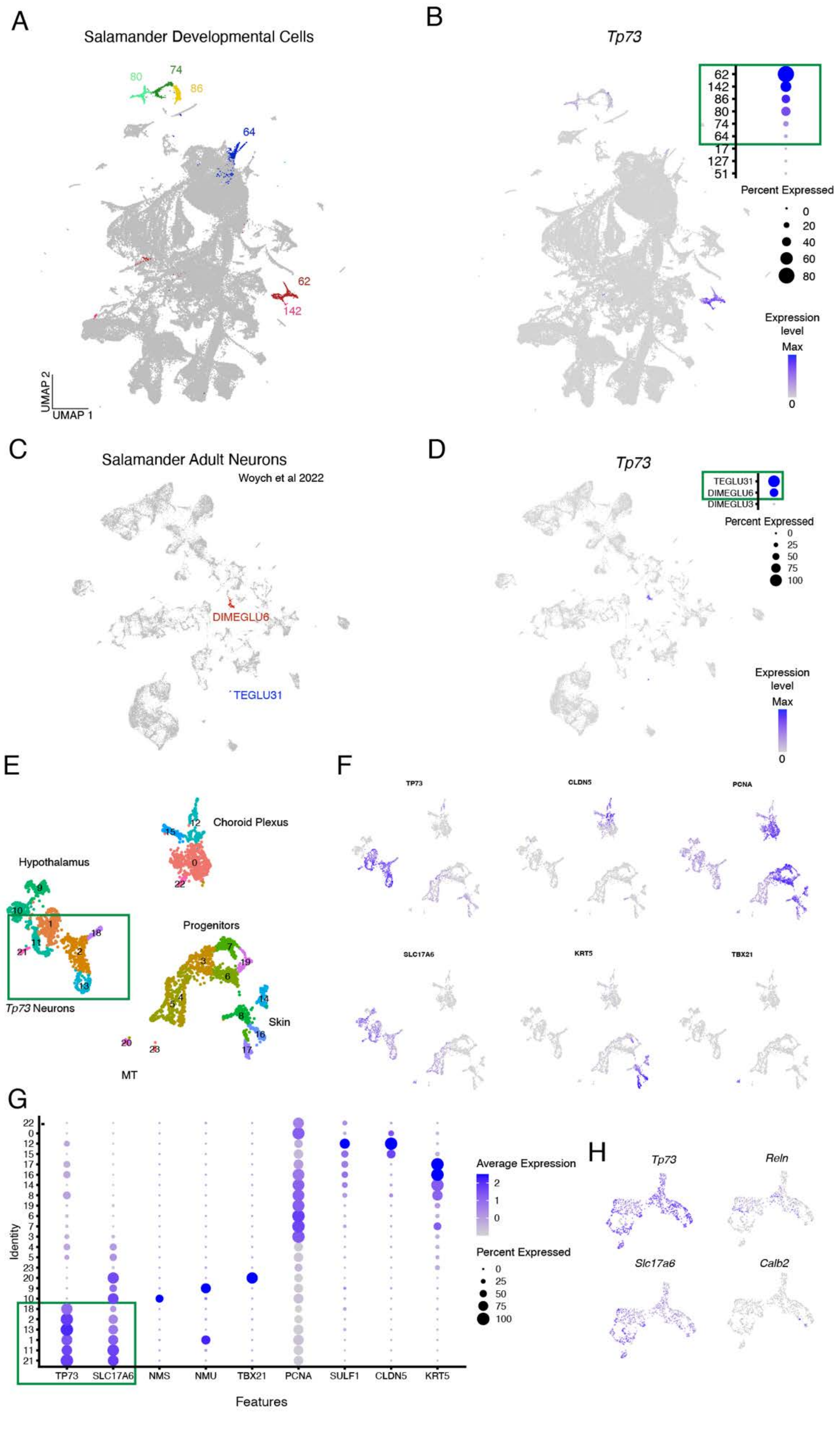
A
Salamander Developmental Cells
80
74
86
64
62
142
UMAP 2
UMAP 1
B
Tp73
62
142
86
80
74
64
17
127
51
Percent Expressed
0
20
40
60
80
Expression level
Max
0
C
Salamander Adult Neurons
Woych et al 2022
DIMEGLU6
TEGLU31
D
Tp73
TEGLU31
DIMEGLU6
DIMEGLU3
Percent Expressed
0
25
50
75
100
Expression level
Max
0
E
Choroid Plexus
Hypothalamus
Progenitors
Tp73 Neurons
Skin
MT
F
TP73
CLDN5
PCNA
SLC17A6
KRT5
TBX21
G
Identity
TP73
SLC17A6
NMS
NMU
TBX21
PCNA
SULF1
CLDN5
KRT5
Features
Average Expression
2
1
0
Percent Expressed
0
25
50
75
100
H
Tp73
Reln
Slc17a6
Calb2

1360

**Supplementary Figure 6. Subsetting of salamander *Tp73* neurons.**
(A-B) UMAP of the full Deryckere et al [40] salamander developmental dataset showing clusters with *Tp73* expression (A). Clusters in dotplot (top left B) ranked by average scaled expression of *Tp73*. Highlighted clusters (colored clusters in (A) and green box in (B)) express *Tp73* in at least 5% of their cells with average scale expression above 0.1, and were selected for further analysis. (C-D) Clusters expressing *Tp73* in adult salamander forebrain dataset [46]. Clusters in dotplot (top left in D) ranked by average scaled expression. Highlighted clusters (colored clusters in (C) and green box in (D)) express *Tp73* in at least 5% of their cells with average scale expression above 0.1, and were selected for further analysis. (E-G) Merge of clusters selected from A-D. Marker genes are plotted (UMAPs in (F), dotplots in (G)) to distinguish *Tp73*+ neurons from other cell types, such as choroid plexus cells (expressing the marker *Cldn5*), proliferating progenitors (*Pcna*+) and skin cells (*Krt5*+). Clusters with high *Tp73* expression (1, 2, 11, 13, 18, and 21, green box) selected for the final *Tp73*+ neuron object. (H) Features plots of *Tp73*+ neurons showing robust *Tp73* and *Slc17a6* expression but scattered *Reln* and mostly absent *Calb2* expression.

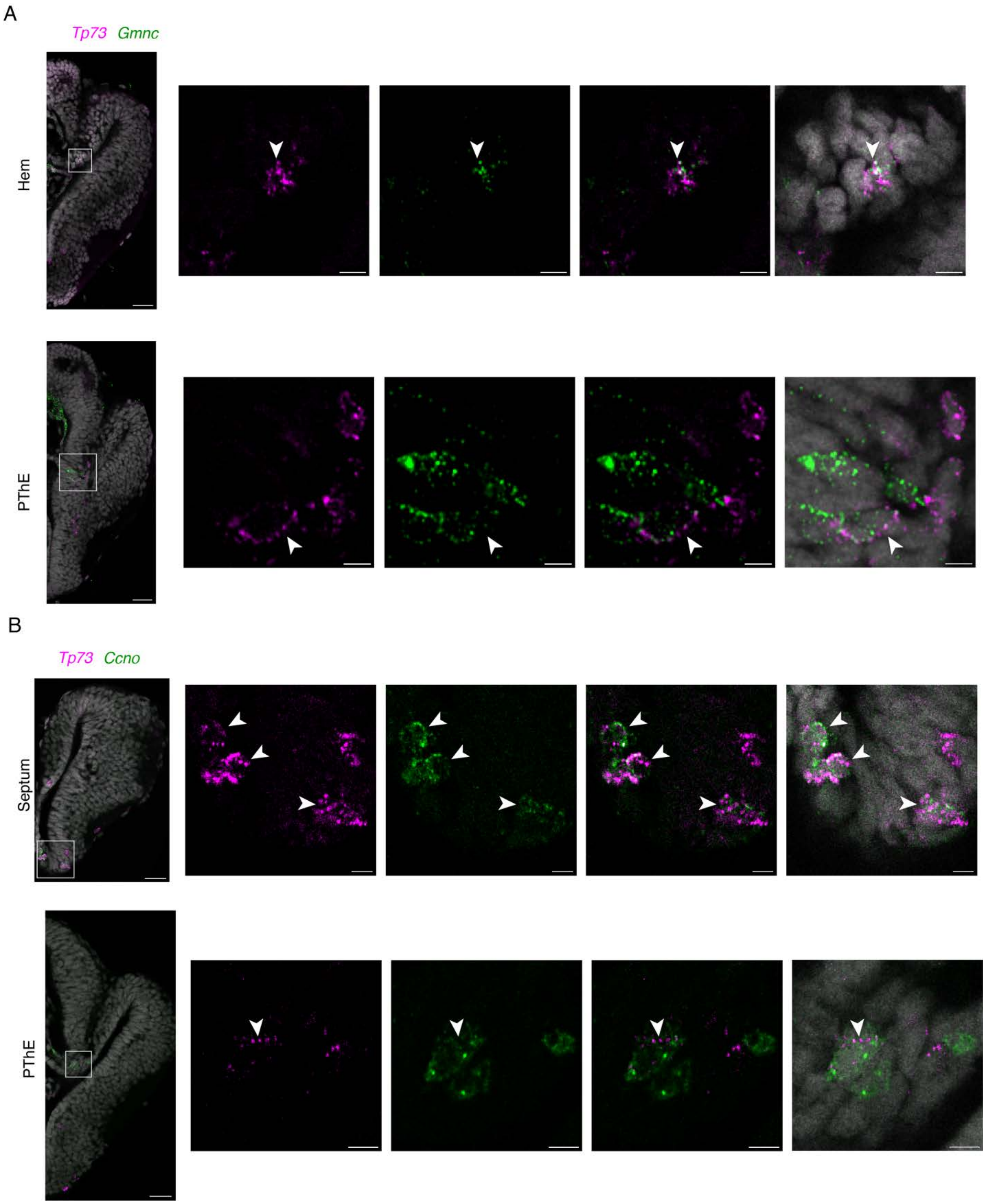


**Supplementary Figure 7. Expression of early multiciliation pathway genes in the developing salamander forebrain.**
(A) Coronal section of stage 41 larval salamander brain showing coexpression of *Tp73* (magenta) and *Gmnc* (green) in hem and PThE. (B) Coronal section of stage 41 larval

salamander brain showing coexpression of *Tp73* (magenta) and *Ccno* (green) in septum and PThE. Scale bars: left 50um, right 10um.

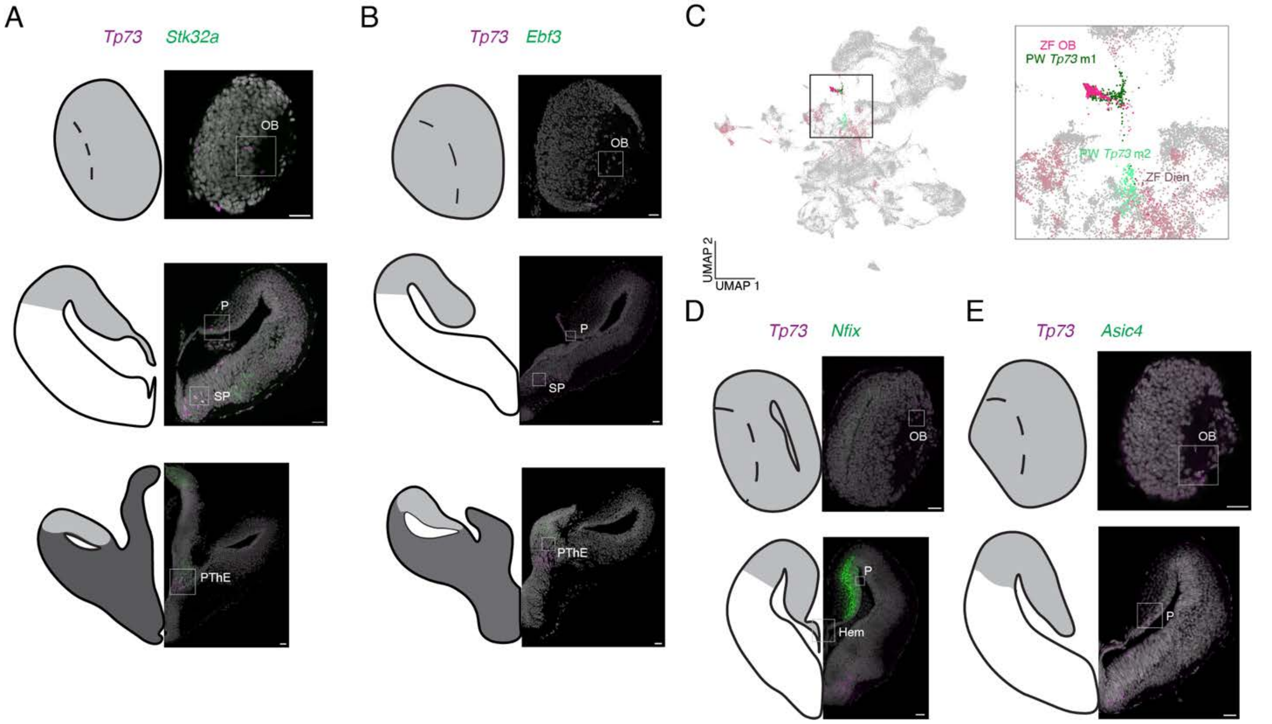


**Supplementary Figure 8. Transcriptomic heterogeneity of salamander *Tp73*+ neurons.**
(A-B) Coronal sections of salamander late active larvae stained for *Tp73* (magenta) and *Stk32a* (A) or *Ebf3* (B). Insets (white boxes) are shown in Fig. 3. Scale bars 50um. (C) Zebrafish and salamander integration (see Fig. 2), showing that zebrafish OB cells co-cluster with salamander m1 class neurons (*Tp73*+ *Ebf3*+), and zebrafish diencephalic cells with salamander m2 class neurons (*Tp73*+ *Ebf3*-) . (F) Coronal sections of salamander larvae stained for *Tp73* (magenta) and *Nfix* (D) or *Asic4* (E). Insets (white boxes) are shown in Fig. 3. Scale bars 50 um.

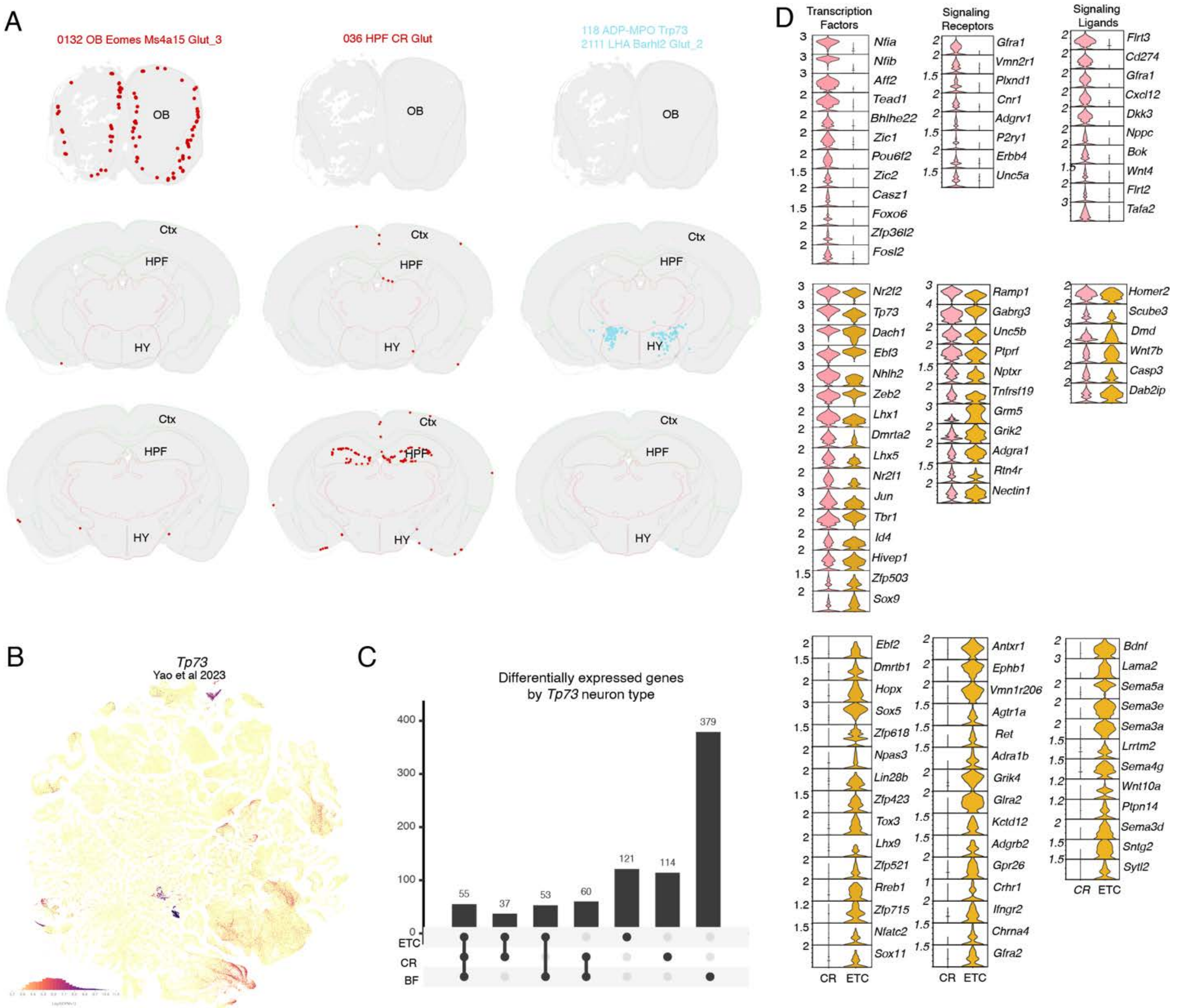


**Supplementary Figure 9. *Tp73*+ neurons in the adult mouse brain and further comparison of mouse CR and ETCs**

(A) Spatial transcriptomic data from Allen Brain Cell atlas [39]. Left: *Tp73*+ ETCs found in OB but not cortex or hippocampus. Middle: CR cells found in cortex and hippocampus but not OB. Right: spatial distribution of basal forebrain *Tp73*+ cells. (B) Feature plot of *Tp73* expression in Allen Brain Cells atlas [39]. (C) Plot showing the number of marker genes that distinguish mouse CR, ETCs or BF cells when compared to the rest of mouse olfactory bulb mitral and tufted cells (see Methods). (D) Violin plots showing expression of marker genes that are shared between ETC and CR or cell-type specific (from the analysis in (C)), only genes annotated with GO terms for transcription factor, signaling receptor, or signaling ligand are shown.

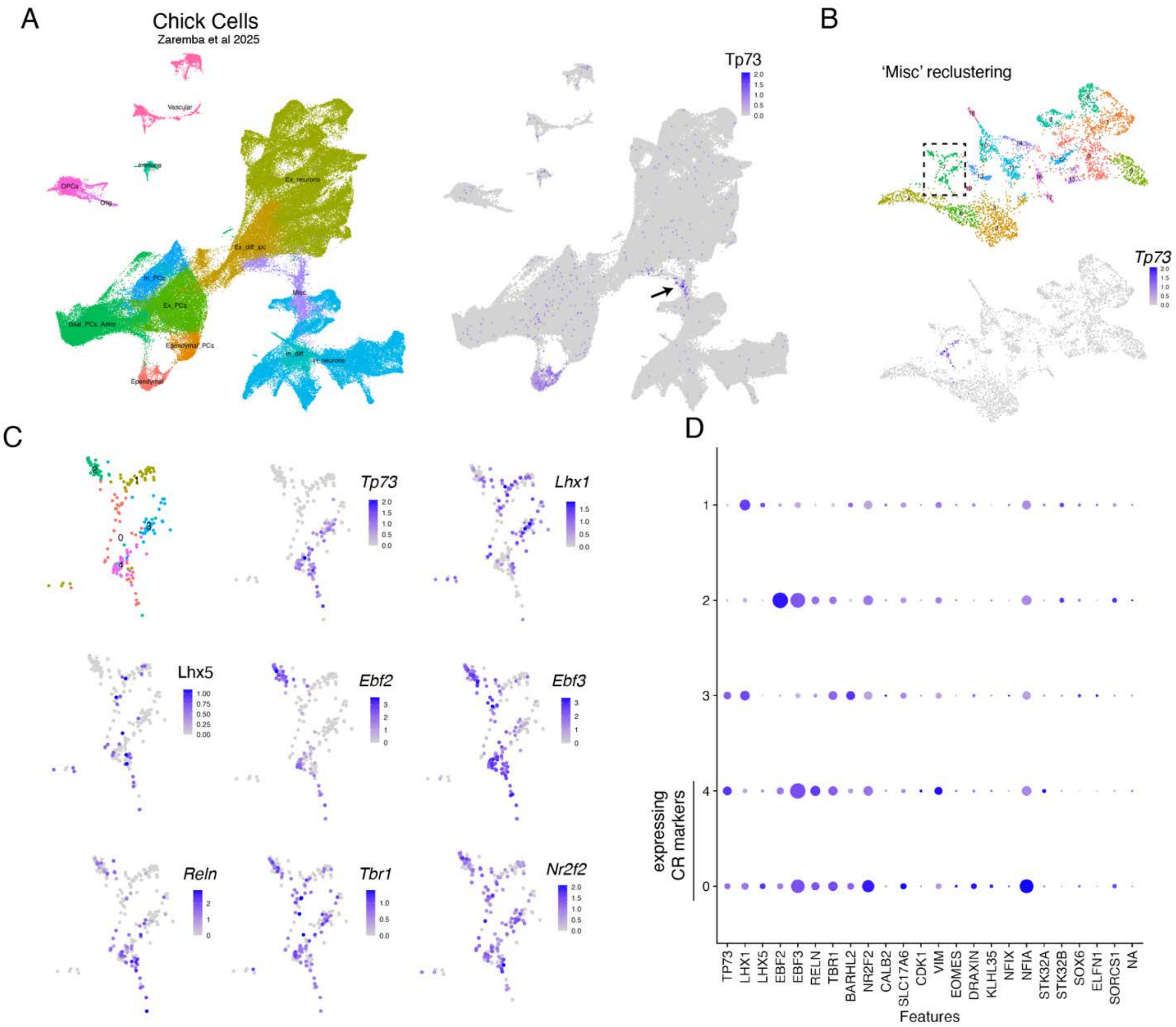


**Supplementary Figure 10 . Expression of CR marker genes in a developing chick scRNA-seq dataset.**

(A) Left: UMAP of all cells from the Zaremba et al 2025 [54] scRNAseq dataset from the developing chick pallium, colored by major class. Right: *Tp73* is sparsely expressed in a neuron class annotated as "misc". (B) Reclustering of misc neuron class reveals one cluster with partial *Tp73* expression. (C-D) Reclustering of the cluster with partial *Tp73* expression shows further heterogeneity; 2 subclusters (0 and 4) express *Tp73* and other known marker genes for CR cells. Data shown as UMAPs (C) and dotplots (D).

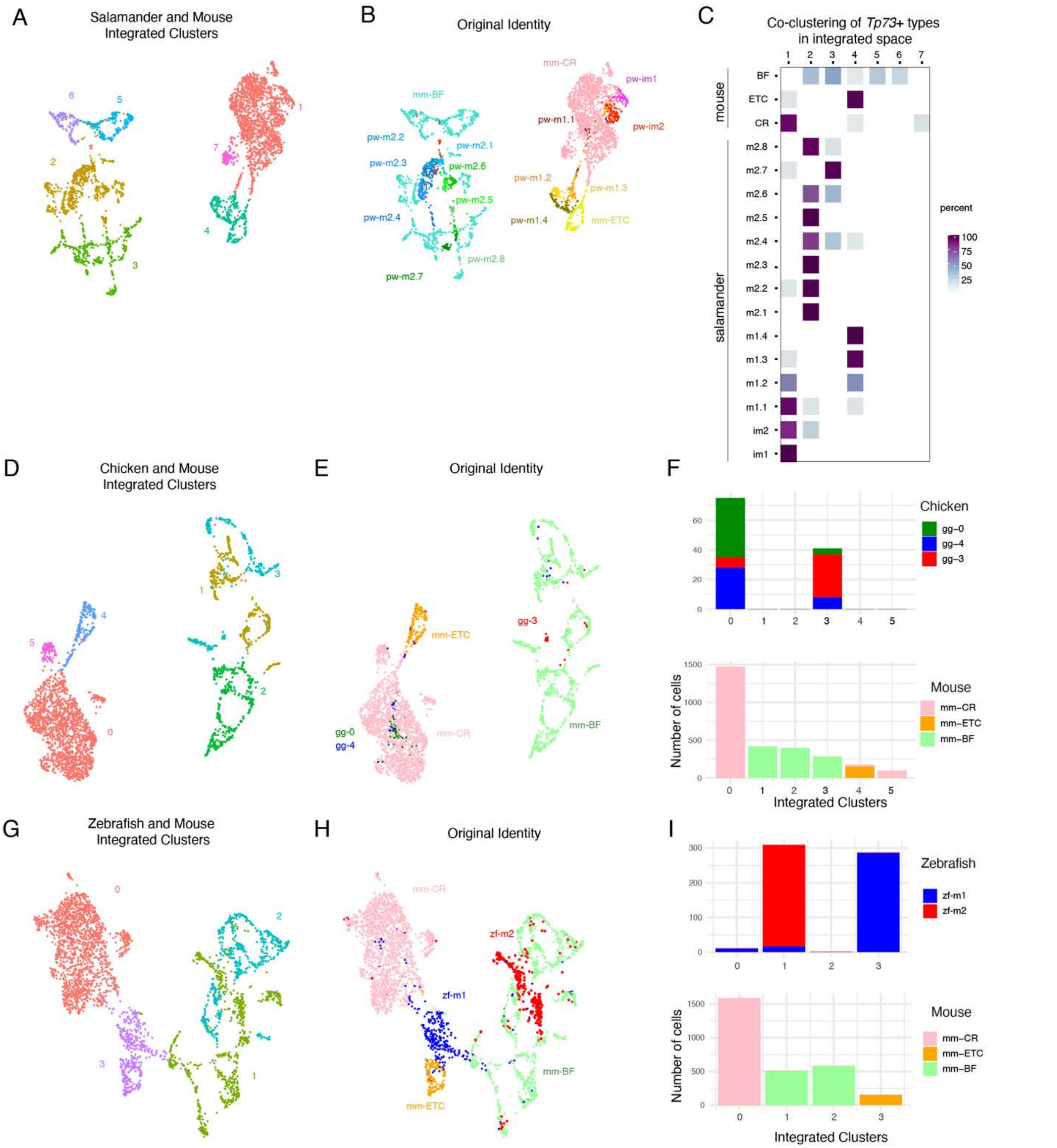


**Supplementary Figure 11. Integration of *Tp73* neurons from chicken and zebrafish with mouse.**

(A) Integrated clusters of chicken *Tp73* neurons from salamander with mouse[39]. (B) The original identity of these integrated neurons as described in Figure 4 (mouse) Figure 3 (salamander) (C) Matrix showing the percentage of cells of each *Tp73* type (original annotations, rows) included the new integrated clusters (columns). (D) Integrated clusters of chicken *Tp73* neurons from Zaremba et al.[54] with mouse[39]. (E) The original identity of these integrated neurons as described in Figure 4 (mouse) and fig. S10 (chicken). (F) Barplot describing composition of each integrated cluster. (G) integrated clusters of zebrafish *tp73* family neurons from Pandey et al.[44] with mouse [39]. (H) The original identity of these integrated neurons. Zebrafish neurons classified based on forebrain integration with salamander (Figure S8C). (I) Barplot describing composition of each integrated cluster.

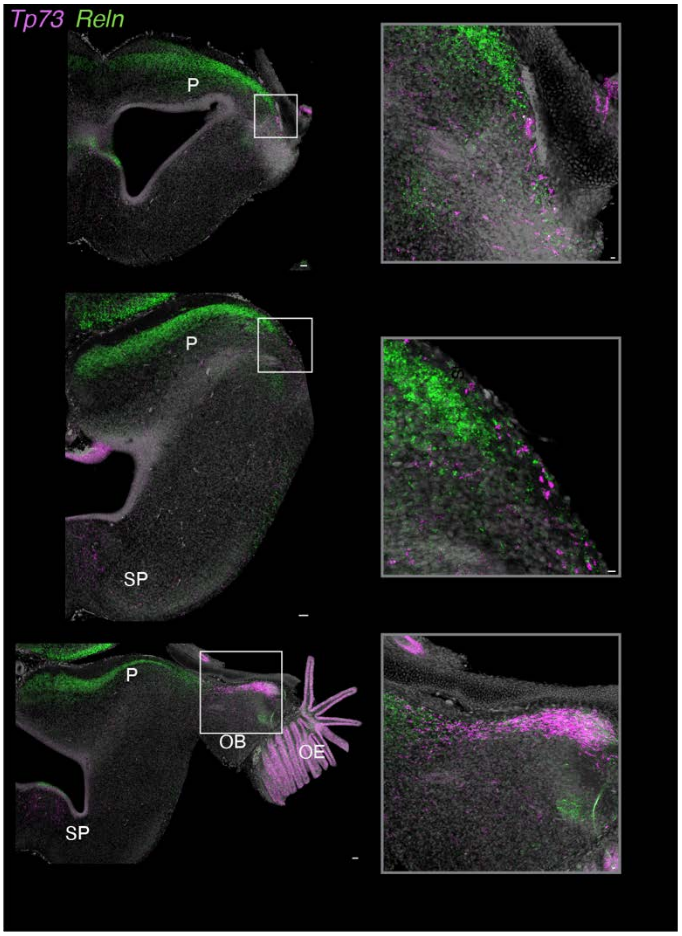


**Supplementary Figure 12. Additional characterization of *Tp73* and *Reln* expression in the developing skate.**
Coronal sections of stage 32 skate brain from anterior (top) to posterior (bottom) showing *Tp73* and *Reln* fluorescent HCR signal (magenta and green, respectively). Strong *Reln* signal was observed in pallial regions, however these *Reln*+ neurons were *Tp73*-negative. Scale bars left 50um, right 10um.

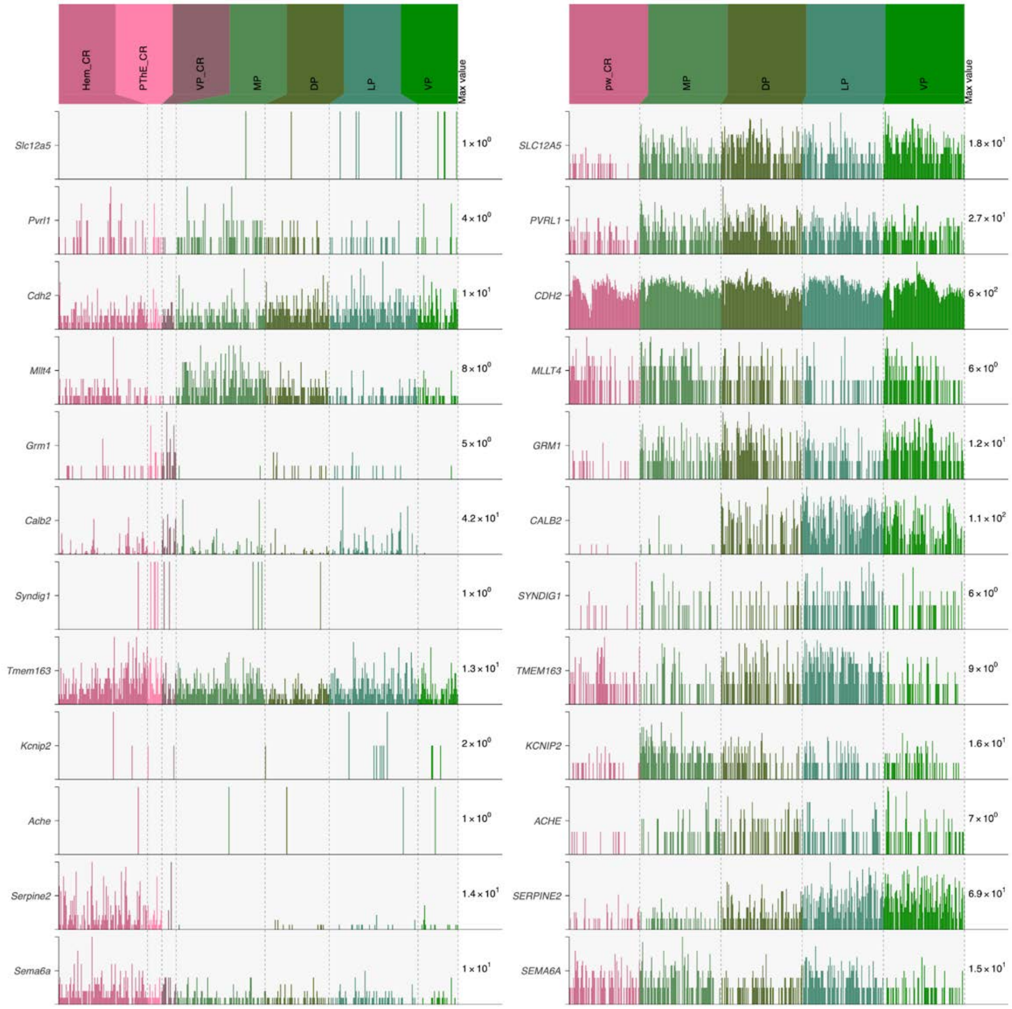


**Supplementary Figure 13. Expression of an additional set of CR marker genes in salamander and mouse.**
Selection of genes known to be positive or negative markers of CR identity in mammals [5,58]. The left panel shows expression in mouse CR cells and pallial glutamatergic neurons [29,33]. The right panel shows expression in mouse salamander cells and pallial glutamatergic neurons.

**Supplementary Video 1**. **Distribution of *Tp73*+ neurons in St46 salamander**
Coronal scan of wholemount stage 46 larvae salamander brain. Labeled with Fluorescent HCR in-situ of *Tp73* (magenta) and *Gad2* (green).

**Supplementary Video 2**. **Distribution of *Tp73*+ neurons in adult salamander**
Coronal scan of wholemount adult salamander brain. Labeled with Fluorescent HCR in-situ of *Tp73* (magenta).

**Supplementary Video 3**. **Distribution of *Tp73*+ neurons in 15 dpf larval zebrafish brain**
Coronal scan of wholemount 15dpf larval zebrafish brain. Labeled with Fluorescent HCR in-situ of *Tp73* (magenta).
